# Mammary mechanobiology: mechanically-activated ion channels in lactation and involution

**DOI:** 10.1101/649038

**Authors:** Teneale A. Stewart, Katherine Hughes, Alexander J. Stevenson, Natascia Marino, Adler L. Ju, Michael Morehead, Felicity M. Davis

## Abstract

A mother’s ability to produce a nutritionally-complete neonatal food source has provided a powerful evolutionary advantage to mammals. Milk production by secretory mammary epithelial cells is adaptive, its release is exquisitely timed and its own glandular stagnation with the permanent cessation of suckling triggers the programmed cell death and tissue remodeling that enables female mammals to nurse successive progeny. Both chemical and mechanical signals control epithelial expansion, function and remodeling. Despite this duality of input, however, the nature and function of mechanical forces in the mammary gland remain unknown. Here, we characterize the mammary force landscape and the capacity of luminal and basal epithelial cells to experience and exert force. We explore the molecular instruments for force-sensing in the mammary gland and the physiological requirement for PIEZO1 in lactation and involution. Our study supports the existence of a multifaceted system of chemical and mechanical sensing in the mammary gland, and a protective redundancy that ensures continued lactational competence and offspring survival.

## Introduction

The adult mammary epithelium consists of an arborized ductal network embedded within an adipose stroma *(1)*. The ductal epithelium contains both luminal and basal cells, which are produced and maintained postnatally by lineage-restricted precursors *(2–7)*. During gestation, a coordinated program of epithelial proliferation, side-branching, differentiation and tissue remodeling takes place *(1, 8)*, resulting in the generation of thousands of alveolar structures that adorn the central ductal epithelium in lobular clusters *(9)*.

The mammary alveolus is the functional unit of the lactating gland. Within this structure, luminal epithelial cells produce and secrete milk into a central lumen and surrounding basal epithelial cells contract in response to maternal oxytocin, expelling milk for the suckling neonate *(10–12)*. At the end of lactation, alveolar mammary epithelial cells are removed in one of the largest physiological cell death cascades that occurs postnatally in mammals *(13, 14)*. This returns the mammary epithelium back to a simple ductal tree, enabling further cycles of regeneration, maturation and milk production over the female’s reproductive lifespan *(15)*.

The formation, function and fate of the adult mammary epithelium is regulated by a range of local and systemic ligands and their receptors, including progesterone, prolactin, oxytocin and leukemia inhibitory factor (LIF) *(16–18)*. In addition to the chemical milieu, mammary epithelial cells reside within a complex physical environment, which may regulate tissue condition and function. Mechanical stresses arising from cell-intrinsic forces (e.g., contractile forces generated by the actin-myosin skeleton) *(12, 19)*, cell-extrinsic forces (e.g., epithelial stretching arising from elevated intralumenal pressure) *(20)* and substrate mechanics (e.g., breast density and tissue tension) *(21, 22)* are thought to affect cell signaling and function in the mammary epithelium. However, whilst roles for mammary mechanotransduction have been proposed *(23, 24)*, the magnitude, direction and dynamics of these forces; their mechanisms of reception and transduction; and their physiological roles and redundancies have not yet been elucidated.

Recently, PIEZO channels have been identified as *bona fide* mechanically-activated ion channels with important roles in mammalian sensory perception *(25)*. In non-neuronal tissue, PIEZO1 has been shown to regulate numerous physiological processes ranging from the maintenance of epithelial barrier integrity *(24, 26)* to lymphatic valve development *(27)*, red blood cell volume *(28)* and myotube formation *(29)*. In the pancreas, PIEZO1 mediates pressure-induced pancreatitis *(30)*, however, roles for PIEZO1 channels in other exocrine organs, including the mammary gland, remain unexplored. Here, we characterize the mammary force landscape and investigate roles for PIEZO1 channels in sensing and transducing epithelial cell forces to sustain or suspend lactation.

## Results

### Luminal and basal mammary epithelial cells experience repetitive, stochastic forces during lactation

The functionally-mature mammary epithelium consists of an inner layer of luminal milk-producing cells and an outer layer of basal milk-ejecting cells (Fig. 1A). Differentiated basal cells express smooth muscle contractile proteins, including α-smooth muscle actin (SMA) *(31)*, and contract in response to maternal oxytocin *(12)*. To measure the magnitude of basal cell contractions, we engineered mice expressing the red fluorescent protein TdTomato in cytokeratin 5 (K5)-positive basal cells (*TdTomato;K5CreERT2* mice). Using 3D time-lapse imaging of intact mammary tissue pieces from lactating reporter mice, we were able to visualize live basal cells and their thin cellular processes *in situ* at high cellular resolution (Fig. 1B). Quantitative assessment of cell morphology before and after oxytocin stimulation revealed a 32.1 ± 0.8% (*P* < 0.05) decrease in surface area during basal cell contraction (Fig. 1B), making these epithelial cell contractions comparable in magnitude to those of cardiomyocytes and intestinal smooth muscle cells *(32, 33)*. To visualize how basal cell-generated forces deform alveolar units for milk expulsion, we loaded mammary tissue from lactating mice with the fluorescent cell-permeable dye CellTracker™ Red, under conditions that led to the preferential labelling of alveolar luminal cells. These data show substantial stochastic deformations to alveolar structures as a consequence of repetitive basal cell contractions with oxytocin stimulation (Supplementary Movie 1 and Fig. 1C).

**Fig. 1.**
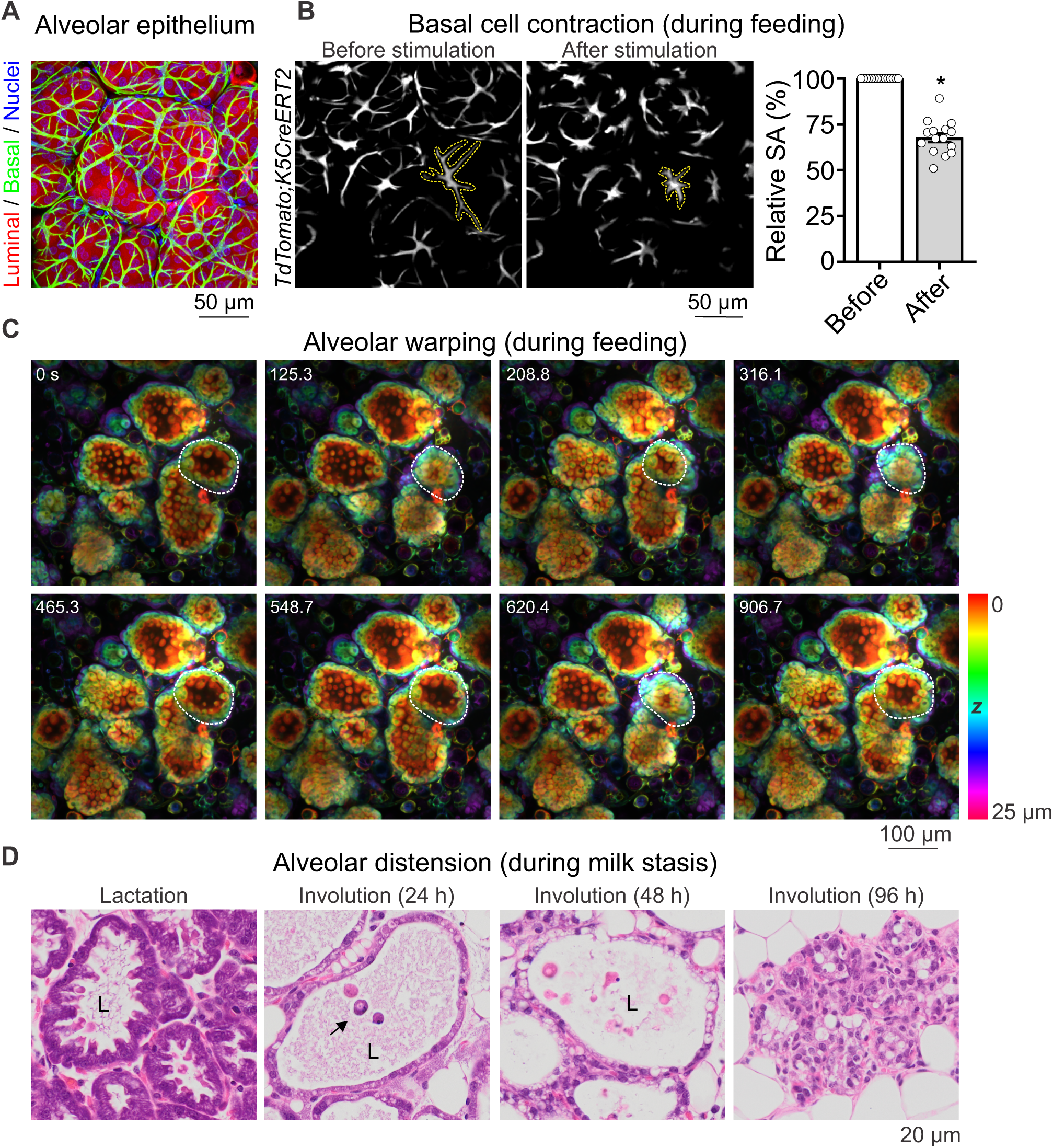
Mammary epithelial cells experience repetitive, stochastic forces during lactation and sustained stretching during early involution. (**A**) 3D maximum intensity projection of lactating mammary tissue showing luminal (secretory) epithelial cells stained with CellTracker™ (red) and basal (contractile) cells with SMA immunostaining (green). Nuclei are blue (DAPI). Representative image from n = 3 mice. (**B**) 4D (*x*, *y*, *z*, *t*) imaging of live lactating mammary tissue from *TdTomato;K5CreERT2* reporter mice, showing basal cell surface area (SA) before and after oxytocin-mediated cell contraction (85 nM). Dotted-line shows a single tracked cell. Graph shows average relative SA before and after oxytocin stimulation (mean ± S.E.M.; 15 cells (total) from n = 3 mice, * *P* < 0.05, student’s t-test). (**C**) 4D imaging of live lactating mammary tissue showing alveolar unit warping due to basal cell-generated force. Tissue was stimulated with oxytocin (85 nM). Dotted line shows a single alveolus through time. Images are depth-coded (see scale bar). See also Supplementary Movie 1. Representative image stack from n = 3 mice. (**D**) Hematoxylin and eosin (H&E) staining of mammary tissue during lactation and involution (24, 48 and 96 h after forced weaning). Arrow shows an apically shed cell; L, alveolar lumen. Representative image from n = 3 mice at each developmental stage. Further images are shown in Fig. S1.

### Milk stasis during involution causes sustained epithelial cell stretching

Post-lactational involution occurs in two stages in mice—the first (reversible) phase (0-48 h), where return of the offspring can lead to the resumption of lactation, and the second (irreversible) phase (> 48 h) marked by alveolar collapse, epithelial cell death and adipocyte regeneration (Fig. 1D) *(15, 34, 35)*. The first phase of involution is characterized by milk accumulation in the alveolar lumen, resulting in increased intralumenal pressure (Fig. 1D and Fig. S1). The sustained overextension of the alveolar epithelium during early involution causes apical cell shedding (Fig. 1D, arrow), a phenomenon that may prolong epithelial barrier integrity by limiting cell density *(24)*.

To quantify the extent of luminal cell compression during involution, we performed immunohistochemistry (IHC) on mouse mammary tissue. Using E-cadherin immunostaining (to define the basolateral cell membranes) and the fluorescent lectin WGA (to demarcate the apical membrane), we were able to visualize large cellular protrusions extending from the lateral contact sites deep into the alveolar lumen (Fig. 2A, arrow). Similar apical curvature was observed in lactating human breast tissue (Fig. S2). Apical projections were absent in tissue samples collected during phase 1 involution (Fig. 2A), suggesting that the apical membrane absorbs a substantial component of the pressure generated through milk stasis. Luminal cell length and area were reduced by more than 50% by the end of the first phase of involution (Fig. 2B).

**Fig. 2.**
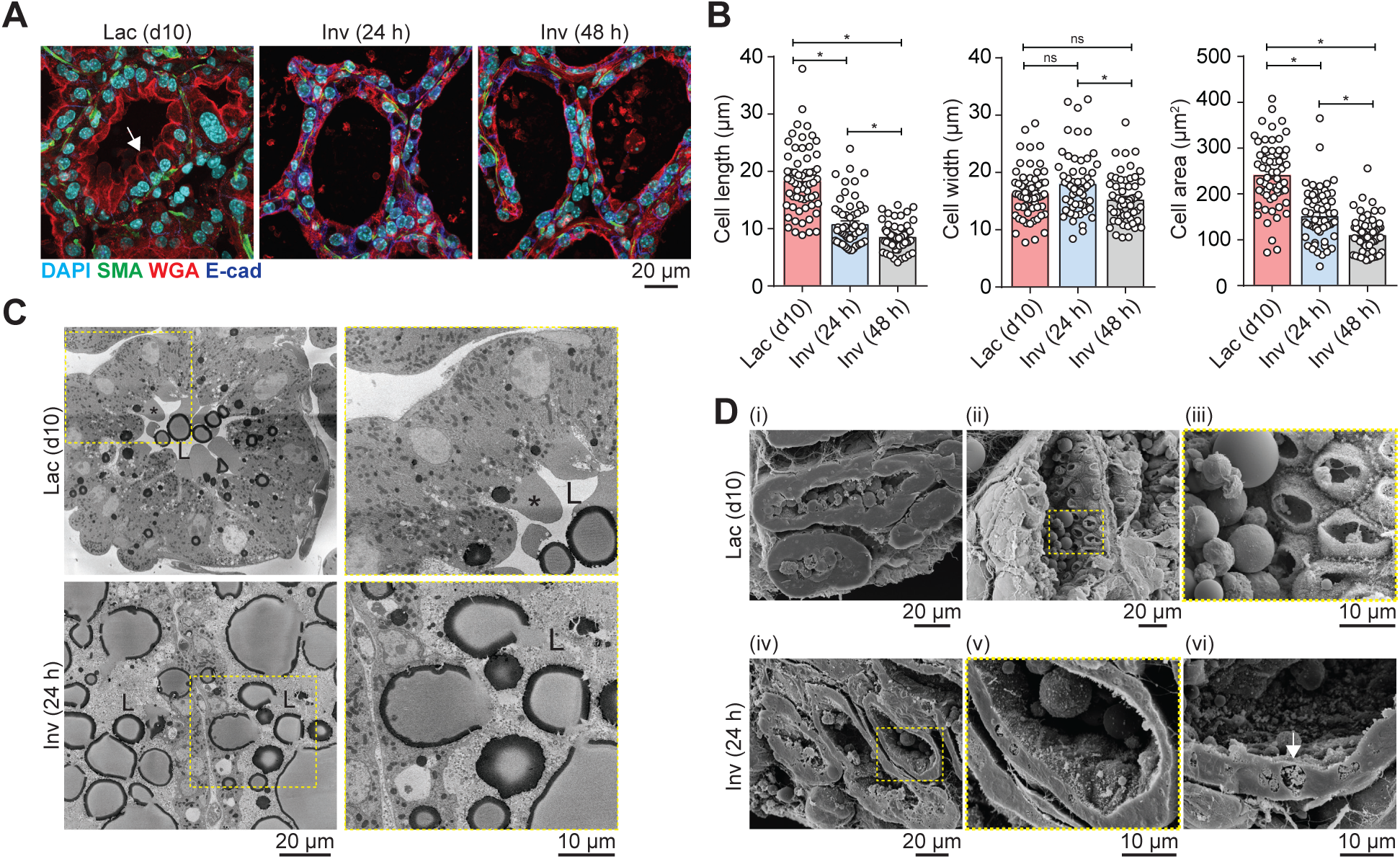
Intralumenal pressure during mammary gland involution is absorbed by the apical cell membrane of luminal epithelial cells. **(A)** Representative immunofluorescence staining of day 10 lactating (Lac d10) and involuting (Inv, 24 and 48 h) mouse mammary tissue. Immunostaining for SMA reveals basal cells (green) and E-cadherin shows the basolateral membranes of luminal epithelial cells (blue). Apical membranes are stained with WGA (red), nuclei are stained with DAPI (cyan). Arrow shows an apical cell protrusion. Representative of n = 3 mice at each developmental stage. **(B)** Quantification of luminal cell length, width and area. Graphs show mean ± S.E.M. from 60 cells from each developmental stage averaged from n = 3 mice, * *P* < 0.05, one-way ANOVA with Bonferroni post-tests; ns: not significant. **(C)** Single slices from SBEM image stacks showing luminal epithelial cells of day 10 lactating (Lac d10) and involuting (Inv 24 h) mouse mammary tissue. Boxed region is magnified in adjacent sub-panel; asterisk shows an apical cell protrusion; L, alveolar lumen. See also Supplementary Movies 2-5. **(D)** SEM imaging of freeze-fractured mouse mammary tissue taken from lactating and involuting (24 h) mice. Boxed region is magnified in next sub-panel; arrow shows endocytic vesicle. Representative images from n = 3 mice. Further images are shown in Figure S4.

We next examined luminal cell morphology and volume using serial block face electron microscopy (SBEM). As expected, apical membrane protrusions were visible during lactation (Fig. 2C, asterisk, and Supplementary Movies 2 and 3), but absent during early involution (Fig. 2C and Supplementary Movies 4 and 5). Interestingly, whilst the plasma membrane calcium ATPase (PMCA) −2 was evenly distributed across the apical membrane of lactating mammary tissue (Fig. S3), protrusions often lacked near-plasmalemmal organelles (Fig. 2C, asterisk and Supplementary Movies 2 and 3). Scanning electron microscopy (SEM) of fractured mammary tissue at different developmental stages enabled visualization of the degree of apical surface distortion during post-lactational involution (Fig. 2D and Fig. S4). Consistent with IHC and SBEM imaging, luminal cell length was reduced during involution (Fig. 2D, (i) vs (iv)). In lactating tissue, microvilli were intact and large secretory vestiges or “cups” were observed (Fig. 2D, (ii) and (iii)). In contrast, apical surfaces were compressed during involution (Fig. 2D, (v) and (vi)) and microvilli could be observed in endocytic vesicles (Fig. 2D, (vi) arrow), consistent with the conversion of these cells to non-professional phagocytes during involution *(13, 36)*.

### Mammary epithelial cells respond to shear stress during lactation

To determine how functionally-mature mammary epithelial cells sense force, we examined intracellular calcium responses in HC11 mouse mammary epithelial cells in a mechanical stimulation assay. HC11 cells treated with lactogenic hormones undergo lactogenic differentiation *(37)*, expressing the lactation markers β-casein (Fig. 3A) and pSTAT5 (Fig. 3B and Fig. S5). Non-differentiated (control) and differentiated HC11 cells were loaded with the ratiometric calcium indicator Fura-5F/AM *(38)* and intracellular calcium responses to fluid shear stress were examined in real-time using a Molecular Devices ImageXpress® Micro high-content imaging system. Using this assay, robotic fluid addition (at a controlled speed and height) inflicts shear stress on HC11 cells cultured on the bottom of the 96-well imaging plate *(39)*. A fraction of non-differentiated HC11 cells responded to shear stress via a transient increase in intracellular calcium (340/380 ratio) (Fig. 3C and Supplementary Movie 6). In contrast, large calcium waves that were propagated from the site of fluid addition were observed in HC11 cells induced to undergo lactogenic differentiation (Fig. 3D and Supplementary Movie 6). These data demonstrate that functionally-mature mammary epithelial cells are able to sense force via a pathway that involves activation of calcium-permeable ion channel(s).

**Fig. 3.**
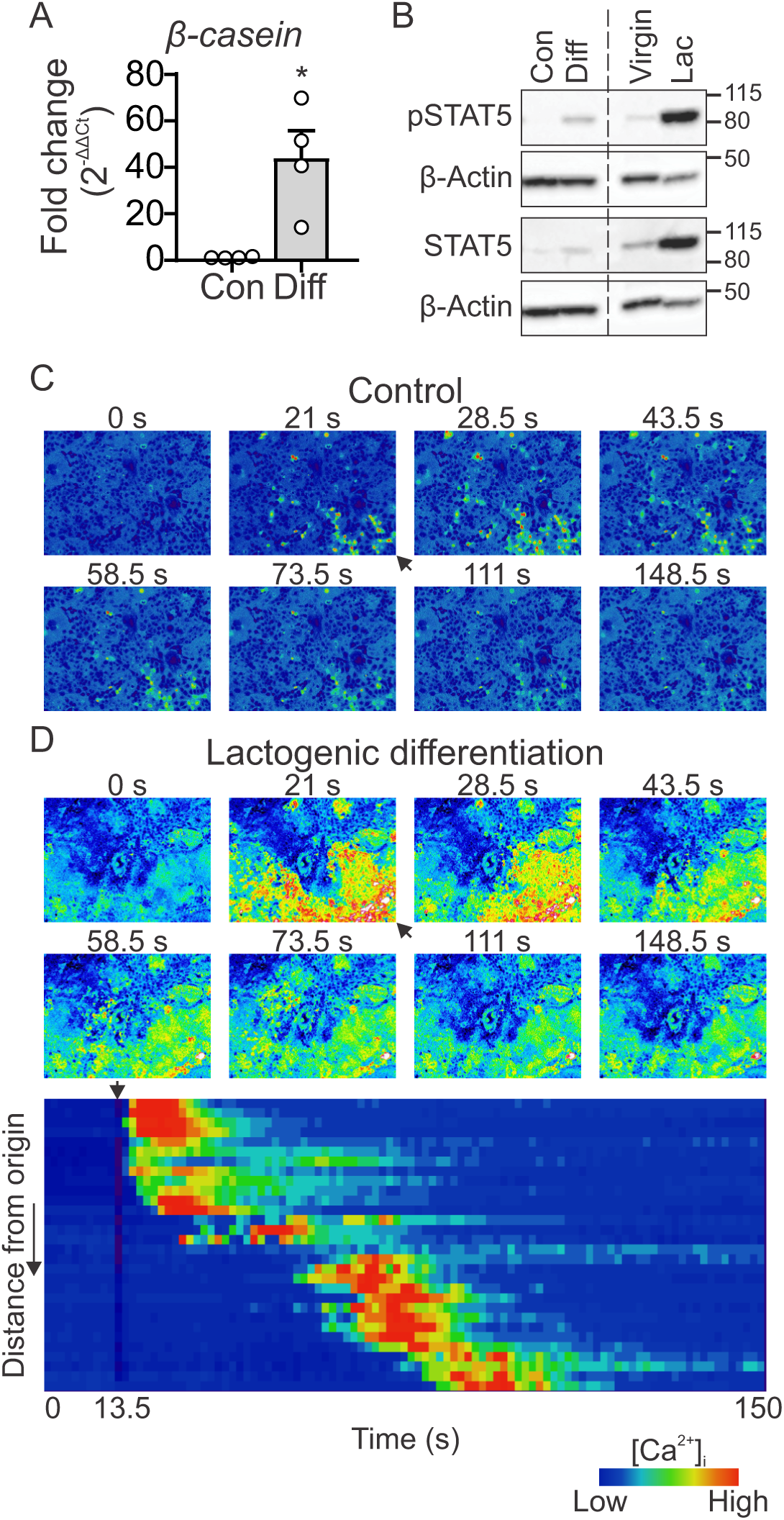
Mammary epithelial cells respond to shear stress in lactation. (**A**) HC11 mouse mammary epithelial cells induced to undergo lactogenic differentiation (Diff) express-casein mRNA. Graph shows mean ± S.E.M., n = 4 independent experiments, * *P* < 0.05, student’s t-test. (**B**) HC11 mouse mammary epithelial cells induced to undergo lactogenic differentiation have higher levels of activated STAT5 (pSTAT5), as observed in protein lysates from lactating mouse mammary tissue (right). Representative immunoblot (n = 4); full immunoblots are shown in Figure S5. (**C**) Non-differentiated (control) HC11 cells stimulated with shear stress (at 15 s) exhibit some transient intracellular calcium responses emanating from the site of fluid addition (black arrow). Images show 340/380 ratio of Fura-5F fluorescence. Representative of 9 wells from n = 3 independent experiments. See also Supplementary Movie 6. (**D**) Differentiated HC11 cells stimulated with shear stress (at 15 s) exhibit a large increase in intracellular calcium propagating from the site of fluid addition (arrow). Images show 340/380 ratio of Fura-5F fluorescence. Heatmap shows relative intensity (F/F_0_, blue to red) of ratioed calcium response as a function of distance from the shear stress origin and time. Representative of 9 wells from n = 3 independent experiments. See also Supplementary Movie 6.

### Functional PIEZO1 is selectively expressed in mature luminal epithelial cells

We performed quantitative RT-PCR to examine the expression of putative and established mechanically-activated ion channels *(40)* in virgin, lactating and involuting mouse mammary tissue (Fig. 4A). RNA samples were validated by *β-casein* mRNA. Although mRNA for the oxytocin receptor (*Oxtr*) is reported to be enriched in purified basal cells during lactation *(41)*, mRNA levels of *Oxtr* and the basal cell marker *Krt14* exhibited a trend to lower expression when examined in lysates prepared from whole tissue (comprised of luminal, basal and stromal cells) (Fig. 4A). This likely reflects the shift in the proportion of basal cells from approximately 33 ± 2.75% of total epithelial cells in virgin ducts to 13 ± 1.38% in lactating alveoli (Fig. S6).

**Fig. 4.**
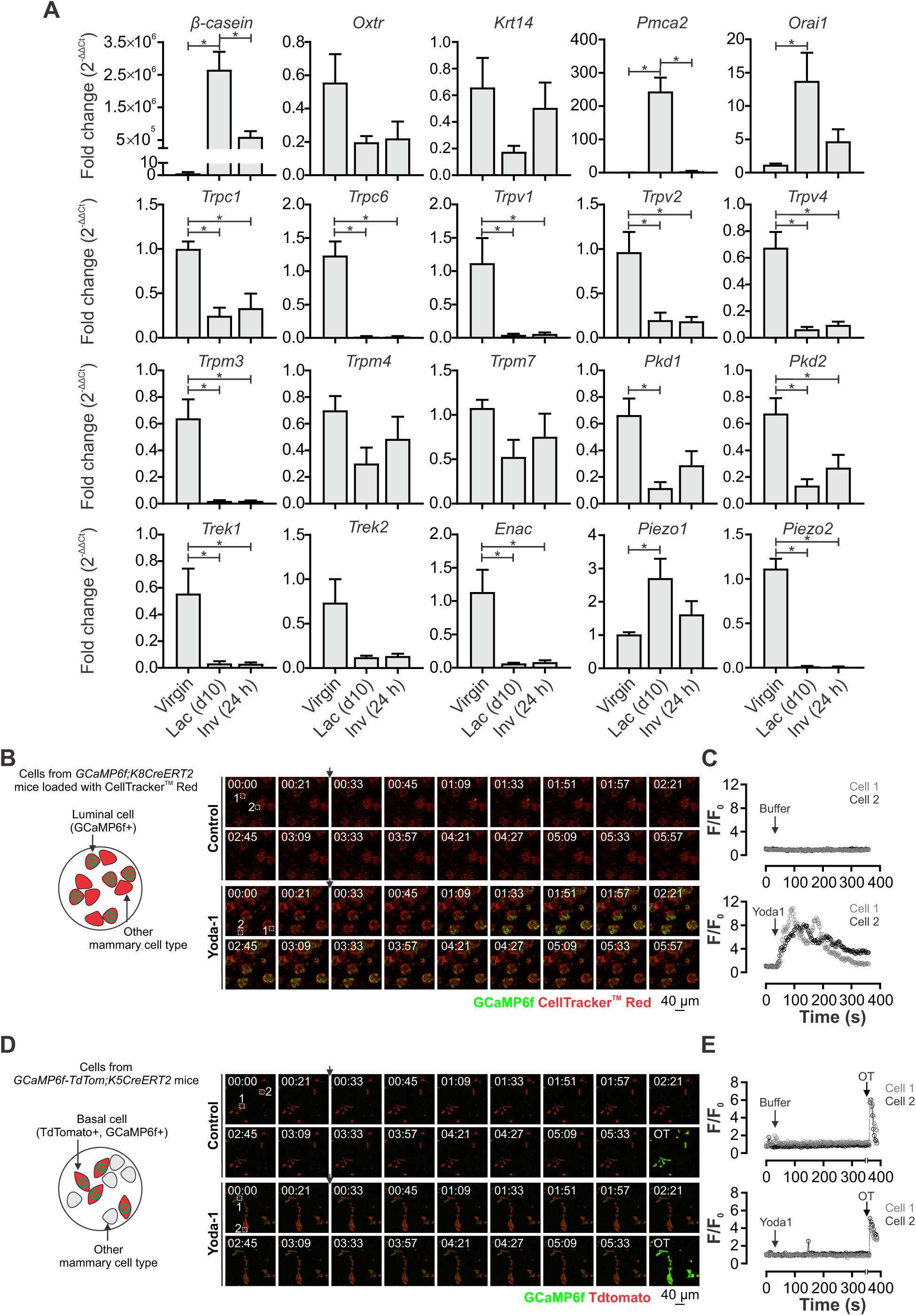
Expression of putative and established mechanically-activated ion channels in lactation and involution. (**A**) Assessment of mRNA levels (qRT-PCR) in whole-tissue lysates isolated from virgin, lactating (day 10) and involuting (24 h) mice. Graphs show mean ± S.E.M., n = 4 mice, * *P* < 0.05, one-way ANOVA with Bonferroni post-tests. mRNA for *Trpa1* and *Traak* were undetectable. (**B**) Intracellular calcium response to the PIEZO1 channel activator Yoda1 in luminal epithelial cells isolated from pregnant *GCaMP6f;K8CreERT2* mice. Cells were loaded with CellTracker™ (red) for morphology, however, only luminal cells in this mixed mammary cell preparation express the calcium sensor (green). See also Supplementary Movies 7 and 8. (**C**) Individual cell traces from two representative cells from each control (buffer only) and Yoda1 - stimulated primary luminal cells. (**D**) Intracellular calcium response to the PIEZO1 channel activator Yoda1 in basal epithelial cells isolated from pregnant *GCaMP6f-TdTom;K5CreERT2* mice. In this mixed cell preparation, basal cells express both TdTomato (red) and the calcium sensor GCaMP6f (green). Cells were stimulated with oxytocin at the end of the assay to demonstrate their ability to respond. See also Supplementary Movies 9 and 10. (**E**) Individual cell traces from two representative cells from each control (buffer only) and Yoda1-stimulated primary basal cells.

As previously reported *(42, 43)*, mRNA levels of the calcium ATPase *Pmca2* and the store-operated calcium entry subunit *Orai1* were significantly enriched in lactating mammary tissue (Fig. 4A). In contrast, gene expression of all transient receptor potential (TRP) calcium-permeable channels analyzed in this study; the potassium channels *Trek1* and *Trek2*; and the epithelial sodium channel *Enac* were unchanged or were significantly reduced in lactating samples (Fig. 4A). *Trpa1* and *Traak* transcripts were undetectable in mammary tissue. Reduced expression of ion channels involved in mechanosensory transduction may be a feature of lactogenic differentiation or could be in-part attributable to their lineage-biased expression (as observed with *Oxtr* and *Krt14*). *Piezo1* was the only mechanically-activated ion channel examined in this study to exhibit increased expression (2.7 ± 0.58-fold, *P* < 0.05) in mammary tissue during lactation (Fig. 4A).

To determine whether PIEZO1 is expressed in both luminal and basal mammary epithelial cell populations, we assessed single cell calcium responses to the selective PIEZO1 channel activator Yoda1 *(44)*. Primary cells isolated from pregnant mice that express the genetically-encoded calcium indicator GCaMP6f *(45)* in mammary luminal cells (*GCaMP6f;K8CreERT2* mice) showed large, sustained increases in intracellular calcium in response to Yoda1 stimulation (Fig. 4B and C and Supplementary Movies 7 and 8). In contrast, GCaMP6f-positive basal cells (isolated from *GCaMP6f-TdTom;K5CreERT2* mice) did not exhibit global increases in cytosolic calcium in response to Yoda1 in single cell recordings (Fig. 4 D and E and Supplementary Movies 9 and 10), but responded, as expected, to stimulation with oxytocin (OT). Based on these data, we focused our studies on defining functional roles for PIEZO1 in luminal mammary epithelial cells during lactation and involution.

### Luminal cell expression of Piezo1 is not essential for lactation and involution

To conditionally delete *Piezo1* in luminal epithelial cells, we generated *Piezo1^fl/fl^;WAPCre* mice. In this model, Cre-mediated excision occurs downstream of activation of the whey acidic protein (WAP) gene promoter in luminal cells during pregnancy/lactation (Fig. 5A) *(46)*. Female mice were taken through one full pregnancy-lactation-involution cycle and the consequence of genetic deletion of *Piezo1* on mammary gland function was assessed after a second pregnancy (Fig. 5B), where > 90% knockdown of *Piezo1* mRNA was observed (Fig. 5C). Alveologenesis and secretory activation were not affected in *Piezo1^fl/fl^;WAPCre* mice (Fig. 5D). Pups nursed by *Piezo1^fl/fl^;WAPCre* mothers gained weight normally (Fig. 5E) and were indistinguishable from pups nursed by control mothers (Fig. 5F), indicating that milk production and secretion were unaffected by luminal *Piezo1* knockdown.

**Fig. 5.**
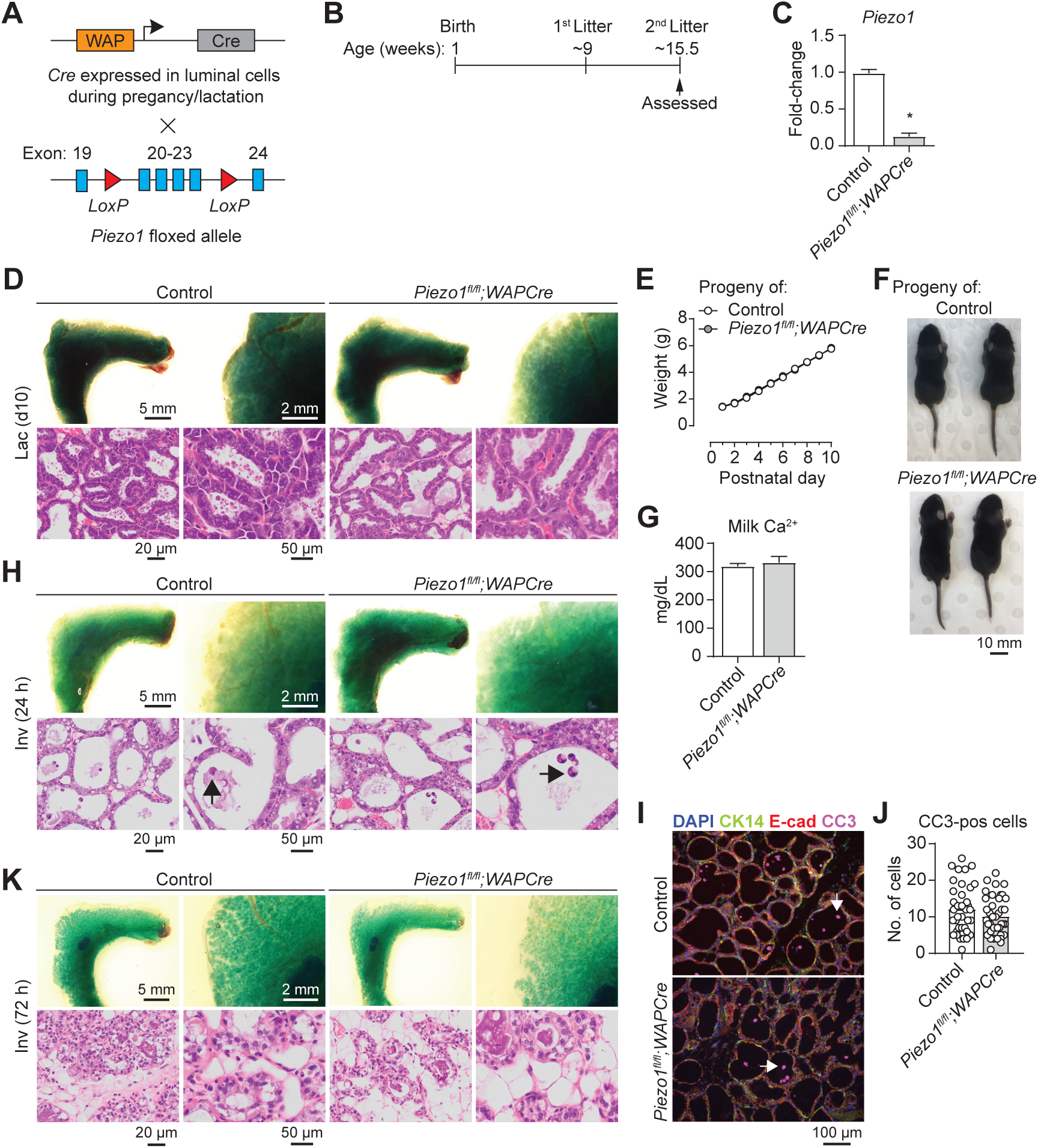
Luminal epithelial cell expression of Piezo1 is not essential for lactation or involution. **(A)** Schematic representation of the *Piezo1* luminal conditional knockout mouse model. **(B)** Experimental timeline for *Piezo1^fl/-fl^;WAPCre* and control mice. **(C)** *Piezo1* mRNA levels in control and experimental *Piezo1^fl/fl^;WAPCre* mice (second lactation). Graph shows mean ± S.E.M., n = 3 mice, * *P* < 0.05, student’s t-test. **(D)** Mammary gland wholemounts (methyl green) and histology (H&E) show normal alveolar development and secretory activation in lactating (d10) control and *Piezo1^fl/fl^;WAPCre* mice (second lactation). Representative of n = 3 mice from each genotype. **(E)** Weight gain in pups nursed by control or *Piezo1^fl/fl^;WAPCre* dams from postnatal day 1 to 10. Graph shows mean ± S.E.M., n = 4 mice. **(F)** Representative images (n = 3 litters per genotype) of pups born to and nursed by control or *Piezo1^fl/-fl^;WAPCre* mice. **(G)** Milk calcium concentrations in control and *Piezo1^fl/fl^;WAPCre* mice. Graph shows mean ± S.E.M., n = 3 mice, *P* > 0.05, student’s t-test. **(H)** Wholemounts (methyl green) and histology (H&E) of control and *Piezo1^fl/fl^;WAPCre* mouse mammary tissue 24 h after forced weaning (second lactation). Representative of n = 4 mice from each genotype. **(I)** Immunofluorescence staining for cleaved caspase 3 (CC3, magenta), E-cadherin (red) and K14 (green) at 24 h involution. Nuclei are stained with DAPI (blue). Representative images from n = 4 mice of each genotype. Arrow shows a CC3 positive apically shed cell. **(J)** Quantification of the number of apically-shed cells in control and *Piezo1^fl/fl^;WAPCre* at 24 h involution. Graph shows mean **±** S.E.M. from 37-40 fields of view from each genotype averaged from n = 4 mice. **(K)** Wholemounts (methyl green) and histology (H&E) of control and *Piezo1^fl/-fl^;WAPCre* mouse mammary tissue 72 h after forced weaning (second lactation). Representative of n = 3 mice from each genotype.

Milk calcium levels were comparable in control and *Piezo1^fl/fl^;WAPCre* mice (Fig. 5G), demonstrating that, unlike store-operated calcium channels *(12)*, mechanically-activated PIEZO1 calcium-permeable channels are not involved in the basolateral flux of calcium that is required for milk calcium enrichment during lactation *(11)*. These data suggest that PIEZO1 channels are either not involved in alveolar morphogenesis and epithelial mechanosensing during lactation, or they have redundant or compensated roles in these processes.

PIEZO1 has been proposed to regulate cell shedding during the early phase of mammary gland involution *(24)*. Gross morphology of mammary glands stained with the histochemical stain methyl green *(47)* were identical in control and *Piezo1^fl/fl^;WAPCre* mice at 24 h involution (Fig. 5H). The number of cleaved caspase-3 (CC3) positive shed cells was also unaffected in *Piezo1^fl/fl^;WAPCre* mice (Fig. 5H-J and Fig. S7) *(48)*. However, histological examination revealed evidence of a precocious involution phenotype in *Piezo1^fl/fl^;WAPCre* mice at 72 h (Fig. 5K). Further studies at later stages of involution are required to determine the potential biological significance of this observation.

### Sensory impulses at the nipple

A combination of visual, auditory and tactile sensory inputs are required for the initiation and maintenance of lactation in most mammalian species *(17)*. Infant suckling activates touch receptors at the nipple, generating signals that travel from the periphery to the central nervous system to trigger oxytocin secretion by hypothalamic neurons *(17)*. Circulating oxytocin is subsequently delivered to the highly-vascularized mammary epithelium, producing the basal cell contractions required for milk ejection and neonate nourishment *(49)*. Our study has revealed that PIEZO1 is not essential for mammary epithelial mechanosensing during lactation (Fig. 5). However, roles for PIEZO channels in other physiological processes that are driven by non-mammary cell types in the mature gland [e.g., in physiological angiogenesis during gestation *(49, 50)* and touch sensation during lactation *(51)*], cannot be discounted. To assess innervation and the capacity for tactile sensation at the nipple during lactation we performed tissue clearing and 3D imaging of mouse nipples (Fig. 6A). Using the neuronal lineage marker PGP9.5, we were able to visualize large nerve bundles in the nipple shaft, base and the glandular tissue adjacent to the primary nipple-draining ducts (Fig. 6B and C). Similar patterns of innervation, adjacent to the milk ducts, were also observed in lactating rabbit teats (Fig. S8). These data reveal an extensive pattern of neural innervation in the mammalian nipple that appears to support the maternal touch response to suckling.

**Fig. 6.**
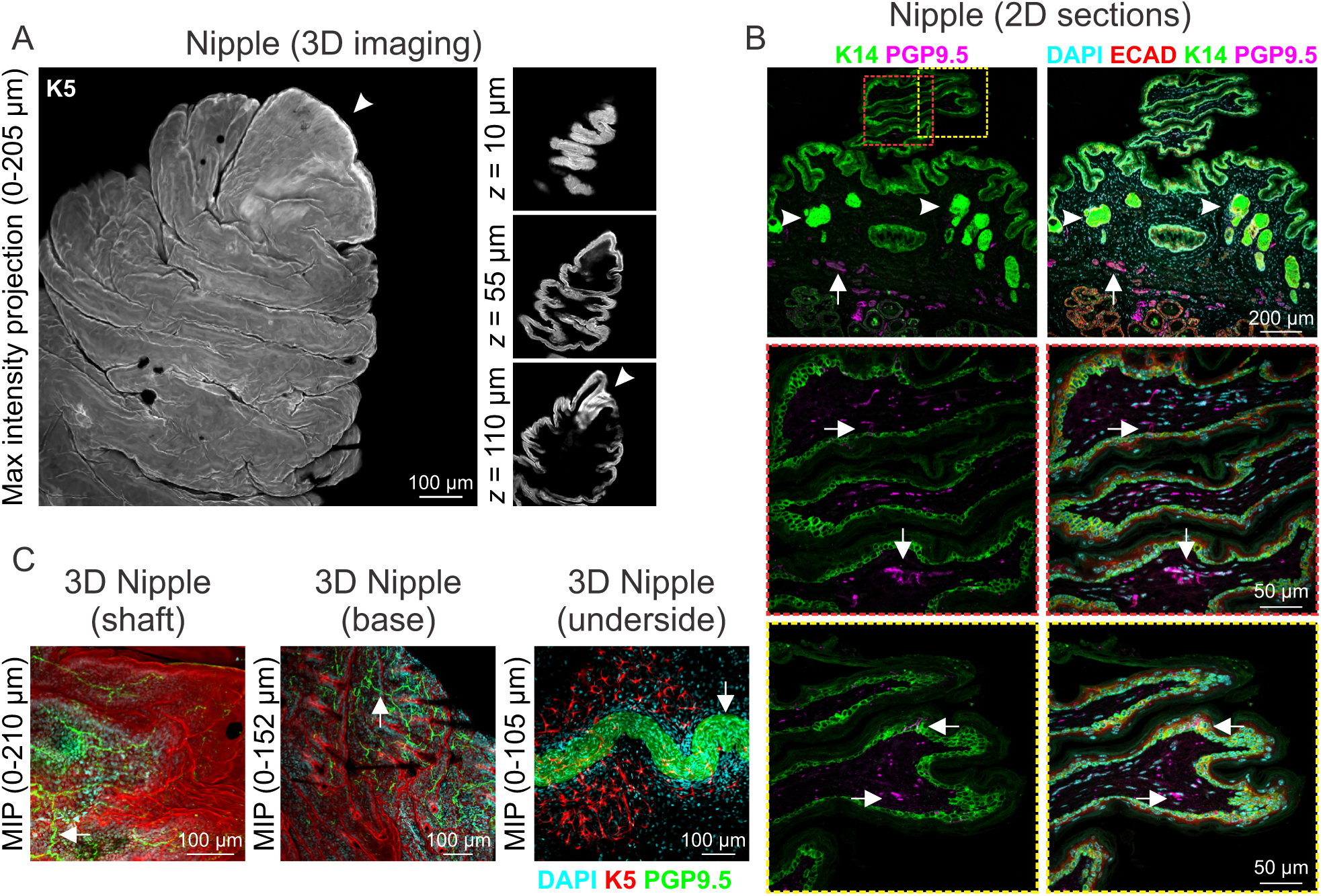
Nipple innervation in the functionally-mature mammary gland. (**A**) 3D maximum intensity projection (*z*: 0-205 m) and optical sections of lactating nipple tissue from *TdTomato;K5CreERT2* reporter mice. (**B**) Immunofluorescence staining for the pan neuronal marker PGP9.5 (magenta). Arrows (top) show PGP9.5 positive staining in nerve bundles at the base of the nipple. Arrows (middle, bottom) show PGP9.5 positive cells in the nipple shaft. Arrow heads (top) show K14 positive sebaceous glands. (**C**) 3D maximum intensity projections [*z*: 0-210 m (left), *z*: 0-152 m (middle), *z*: 0-105 m (right)] of PGP9.5 wholemount immunostaining (green) in cleared nipple tissue from the nipple shaft, base and underside (mammary gland) of *TdTomato;K5CreERT2* reporter mice. Representative images from n = 3 mice.

## Discussion

The perception and transduction of mechanical signals is ubiquitous in nature. Plants alter their growth and physical form in response to a range of environmental force stimuli, including wind, touch and gravity *(52)*. Mechanical inputs guide locomotion in insects *(53)*. In mammals, mechanosensation underpins fundamental biological processes, ranging from the perception of sound and touch *(51, 54)* to the regulation of limb position *(55)* and blood pressure *(56)*. An integrated approach to understanding the function and malfunction of tissues in complex multicellular organisms, which incorporates both mechanical and biochemical elements, is therefore essential.

Here, we characterized the nature and scale of the mechanical load that mammary epithelial cells experience during lactation and the post-lactational period. These data reveal that, similar to contractile cells in other force-generating organs *(32)*, basal cells in the mammary gland undergo substantial radial shape changes during contraction and relaxation. Using 4-dimensional, *ex vivo* tissue imaging, we were able to demonstrate how this basal cell-mediated force is transmitted to the inner layer of compliant luminal epithelial cells. Finally, we characterized a third form of force in the functionally-mature mammary epithelium, which arises due to milk accumulation in the alveolar lumen, and were able to visualize for the first time the physical footprint of this force on the apical membrane of alveolar luminal epithelial cells.

How mechanical forces are sensed by luminal and basal epithelial cells in the namesake organ of mammals remains an outstanding question in mechanobiology. It is also unclear whether these forces direct or drive mammary form and function, or whether the tissue develops, functions and remodels *despite* these mechanical inputs. Our observation that the majority of mammalian mechanically-activated ion channels are downregulated during lactation, supports the latter theory. In our study, PIEZO1 was the only mechanosensitive ion channel to be upregulated during lactation, where stimulation with the chemical activator, Yoda1 *(44)*, produced an intracellular calcium response in the luminal epithelial lineage. These tantalizing data suggest that mammary epithelial cells are able to dynamically-adapt their mechanosensing machinery in a developmental stage specific manner, to “channel” force-sensing in lactation through PIEZO1.

To investigate roles for PIEZO1 in the alveolar epithelium during lactation, we generated a luminal cell-specific *Piezo1* knockout mouse. In this model, lactation and involution proceeded without major impairment. It is currently unclear what role upregulated and activatable PIEZO1 plays in the luminal epithelium, whether another mechanosensitive ion channel is able to alter expression and compensate in its absence, or whether inherent biological redundancies exist. In other cell-systems, e.g., stomatal stem cells of *Arabidopsis*, chemical signals are able override mechanical input to establish cell polarity orientation *(52)*. Similar signal duplication with chemical domination may also exist in mammalian cells and systems. Indeed, remarkable redundancy in the creation of the alveolar epithelium by mammary stem/progenitor cells has already been observed in this evolutionarily essential organ *(3, 7)*.

Using tissue clearing and deep 3D imaging, we revealed the intricate structure of the lactating nipple, where the physical sensation of suckling initiates milk letdown through hypothalamic oxytocin release *(17)*. We postulate that the related PIEZO protein, PIEZO2, mediates touch sensitivity at the nipple, via Merkel cells and/or sensory neurons *(51)*, providing another distinct but related avenue for mechanosensing in lactational physiology. However, redundancies in sensory input (tactile, auditory and visual) are also likely to exist in most mammals *(17, 57)*, enabling the mammary gland to fulfil its sole biological role, which is essential for offspring survival in nearly every species in the class Mammalia. The lactating mammary gland, therefore, truly is a “force” to be reckoned with!

## Materials and Methods

### Mice

All experimentation was carried out in accordance with the *Australian Code for the Care and Use of Animals for Scientific Purposes*, the *Queensland Animal Care and Protection Act (2001)*, the National Institutes of Health’s *Guide for the Care and Use of Laboratory Animals*, the Animal (Scientific Procedures) Act 1986, and the European Union Directive 86/609 with local animal ethics committee approvals. Animals were kept in a Specific Pathogen Free (SPF) facility, and housed in individually-ventilated cages under a 12:12 h light-dark cycle, with water and food available *ad libitum*. The following mice were purchased from The Jackson Laboratory (Bar Harbor, ME): *WAP-Cre* (B6.Cg-Tg(Wap-cre)11738Mam/JKnwJ, stock no. 008735) *(46)*, *Piezo1-flx* (Piezo1^tm2.1Apat^/J, stock no. 029213) *(28)*, *K8-CreERT2* (Tg(Krt8-cre/ERT2)17Blpn/J, stock no. 017947) and *K5-CreERT2* (B6N.129S6(Cg)-Krt5^tm1.1(cre/ERT2)Blh^/J, stock no. 029155). *TdTomato-flx* mice were a kind gift from Prof. Ian Frazer (University of Queensland). *GCaMP6f-flx* mice were a kind gift from Dr. James W. Putney (NIEHS/NIH). C57BL6/J mice were obtained from the Animal Resources Centre (Western Australia). All strains were acquired and maintained on a C57BL6/J background. *WAP-Cre, K8-CreERT2* and *K5-CreERT2* mice were maintained as hemi/heterozygotes. Genotyping was performed on mouse toe, ear or tail DNA by PCR using the following primers: to distinguish Cre positivity in the *WAP-Cre* and *K8-CreERT2* models 5’-GCG GTC TGG CAG TAA AAA CTA TC-3’, 5’-GTG AAA CAG CAT TGC TGT CAC TT-3’ (transgene 100 bp) and 5’-CTA GGC CAC AGA ATT GAA AGA TCT-3’, 5’-GTA GGT GGA AAT TCT AGC ATC ATC C-3’ (internal positive control 324 bp); to distinguish *Piezo1-flx* 5’-GCC TAG ATT CAC CTG GCT TC-3’ and 5’-GCT CTT AAC CAT TGA GCC ATC T-3’(wildtype 188 bp, mutant 380 bp); to distinguish *K5-CreERT2* 5’-GCA AGA CCC TGG TCC TCA C-3’, 5’-GGA GGA AGT CAG AAC CAG GAC-3’, 5’-ACC GGC CTT ATT CCA AGC-3’ (wildtype 322 bp, mutant 190 bp); to distinguish *TdTomato-flx* and *GCaMP6f-flx* 5’-CTC TGC TGC CTC CTG GCT TCT-3’, 5’-CGA GGC GGA TCA CAA GCA ATA-3’ and 5’-TCA ATG GGC GGG GGT CGT T-3’ (wildtype 330 bp, mutant 250 bp).

To induce recombination in *K5CreERT2* and *K8CreERT2* mice, animals were injected with tamoxifen (1.5 mg) in sunflower oil at 4-weeks of age. A further three tamoxifen injections were administered every second day on alternating sides at 8-weeks of age (total dose, 6 mg). The dose and timing of tamoxifen administration (1.5 mg at the onset of puberty) had little effect on ductal morphogenesis *(58)* and resulted in 95.7 ± 1.6% recombination in basal cells. Mice were taken through a 6-week washout before mating. To induce recombination in *WAPCre* mice, animals were taken through one full pregnancy-lactation-involution cycle. Pups (average 8; range 6-10) were weaned on postnatal day 21. Mothers were mated one week later in their second (experimental) mating. For all experimental litters pup number was standardized to 6-8.

To obtain tissue during lactation and involution, adult female mice were mated with C57BL6/J sires and allowed to litter naturally. Lactating tissue was harvested on day 10 of lactation. For forced involution studies, pups were removed on day 10 of lactation (range 10-12) and tissue was harvested from involuting mothers 24, 72, or 96 h later, as indicated.

### Cell lines

HC11 cells (originally isolated from the mammary gland of a pregnant mouse) *(59)* were a gift from Prof Melissa Brown (The University of Queensland, Australia). Cells were maintained in Roswell Park Memorial Institute 1640 (RPMI-1640) medium with L-glutamine and sodium bicarbonate (R8758, Sigma-Aldrich), supplemented with 10% fetal bovine serum (FBS; Thermo Fisher Scientific), bovine insulin (5 µg/mL, I6634, Sigma-Aldrich) and murine epidermal growth factor (10 ng/mL, E4127, Sigma-Aldrich). To induce differentiation, HC11 cells were first cultured for 48 h in maintenance media, followed by 24 h in pre-differentiation media containing RPMI-1640, FBS (10%) and bovine insulin (5 µg/mL). Cells were grown for a further 96 h in differentiation media containing RPMI-1640, FBS (10%), bovine insulin (5 µg/mL), ovine prolactin (5 µg/mL, L6520, Sigma-Aldrich) and dexamethasone (1 µM, D4902, Sigma-Aldrich), with daily media changes. HC11 cells were maintained in a humidified incubator at 37°C with 5% CO_2_, and routinely (every 6 months) tested negative to mycoplasma (MycoAlert, LT07-218, Lonza).

### Human subjects

Breast tissue biopsies from lactating women were obtained from the Susan G. Komen Tissue Bank at the IU Simon Cancer Center, USA *(60)*. All samples were obtained with informed consent and patients (female, aged 25-42) were recruited under a protocol approved by the Indiana University Institutional Review Board (IRB protocol number 1011003097) and according to The Code of Ethics of the World Medical Association (Declaration of Helsinki). The current project received additional, site-specific approval from the Mater Misericordiae Ltd Human Research Ethics Committee. Tissue was formalin fixed and paraffin embedded as per standard protocols. Women classified as lactating had at least 1 live birth (range 1-4) and were actively breastfeeding at the time of tissue donation (at least once daily).

### Live cell imaging

HC11 sheer stress calcium imaging experiments were performed in physiological salt solution (PSS; 10 mM HEPES, 5.9 mM KCl, 1.4 mM MgCl_2_, 1.2 mM NaH_2_PO_4_, 5 mM NaHCO_3_, 140 mM NaCl, 11.5 mM glucose, 1.8 mM CaCl_2_; pH 7.3-7.4). For shear stress experiments, HC11 cells were plated in 96-well, optical-grade, TC-treated, black microplates (353219, BD Biosciences) and differentiated (as indicated), as described above. HC11 cells were loaded with the ratiometric calcium indicator Fura-5F/AM (4 μM, 6616, Setareh Biotech) for 30 min at 37°C as previously described *(12)*. Cells were stimulated by the addition of 100 μL of PSS-Ca^2+^ at a defined rate using an ImageXpress Micro XLS widefield high-content system with fluidic control (Molecular Devices) *(61)*. Images were recorded every 1.5 s for 150 s using a 10× Plan Fluor objective. The ImageJ custom built plugin Ratio Plus *(62)* was used to create 340/380 ratios and fluorescence intensity calculated based on starting fluorescence (F_0_) *(12)*.

For live imaging of primary cells isolated from *GCaMP6f;K8CreERT2* and *GCaMP6f-tdTom;K5CreERT2* mice, tissue was removed and cells dissociated as previously described *(63)*. Following dissociation, cells were left to attach for at least 3 h in a humidified incubator at 37°C with 5% CO_2_. *GCaMP6f;K8CreERT2* primary cells were loaded with CellTracker™ Red (1 µM, C34552, Life Technologies) in DMEM/F12 containing 10% FBS and gentamycin (50 µg/mL) for at least 30 min at 37°C prior to imaging. Primary cells were imaged in 100 µL DMEM/F12 containing 10% FCS and gentamycin (50 µg/mL) and were stimulated with either 150 µL PSS-Ca^2+^ or Yoda1 (20 µM, 5586, Tocris, prepared fresh) in PSS-Ca^2+^ (final concentration of Yoda1 is 12 µM). Images were acquired on an Olympus FV3000 inverted confocal microscope using a UPLSAPO 30×/1.05 silicone objective, resonant scanner and the z-drift compensator (ZDC).

### Ex vivo *4D tissue imaging*

Live 3D time-lapse (4D) imaging was performed as previously described *(12)*. Tissue was bathed in PSS-Ca^2+^ and stimulated with oxytocin (85 nM, O3251, Sigma-Aldrich) in PSS-Ca^2+^.

### Mouse milking

For mouse milking studies, lactating dams were removed from the nest for 2 h prior to milking on lactation day 5. Lactating mice were lightly anesthetized using isoflurane (2%). Mice were administered oxytocin (2 IU) by intraperitoneal injection and milking from the right abdominal mammary gland was initiated 2 min later. Milk was expressed by light manual manipulation of the mammary gland and milk was immediately collected from the tip of the nipple using a pipette.

### Milk Ca^2+^ measurements

Free calcium concentrations in mouse milk were determined using a phenolsulphonephthalein reaction with the QuantiChrom Calcium Assay Kit (DICA-500, BioAssay Systems) as per manufacturer’s instructions and as previously reported in mouse milk samples *(12)*.

### Histology and immunofluorescence staining

Mouse tissue was dissected, spread on biopsy pads and fixed in neutral buffered formalin (NBF, 10%) overnight at room temperature. Routine processing and paraffin embedding was performed; immersion in 70% ethanol (45 min), 90% ethanol (45 min), 95% ethanol (45 min), 100% ethanol (45 min, ×3), xylene (30 min, ×3) and paraffin (1 h, 60°C, ×3). Formalin-fixed paraffin embedded (FFPE) slides were cut at 4-5 μm. Immunofluorescence staining was performed by heat-induced epitope retrieval, as previously described *(3)*. Briefly, FFPE slides were deparaffinized by immersion in xylene (3 × 5 min) and rehydrated through a reducing ethanol series. Permeabilization was performed by immersion in PBS containing Triton X-100 (0.5%). Antigen retrieval in sodium citrate buffer (0.01 M, pH 6) was performed using a Decloaking Chamber NxGen digital pressure system (Biocare Medical) at 110°C for 11 min. Tissue sections were blocked for 1 h in normal goat serum (10%) in PBS containing Triton X-100 (0.05%). Primary antibody incubation was performed overnight in a humidified chamber at 4°C. Secondary antibody incubation (1:500) and wheat germ agglutinin (WGA; 5 μg/mL) staining was performed for 1 h at room temperature. Nuclear DAPI (625 ng/mL) staining was performed for 10 min at room temperature. The following primary antibodies and dilutions were used in this study: rabbit anti-SMA (1:400-800), rabbit anti-E-cadherin (1:400), mouse anti-E-cadherin (1:200), rabbit anti-CC3 (1:200), chicken anti-K14 (1:200), anti-rabbit PGP9.5 (1:100) and rabbit anti-PMCA2 (1:200). The following secondary antibodies were used: goat anti-mouse Alexa Fluor 647 (Life Technologies), goat anti-rabbit Alexa Fluor 647 (Life Technologies), goat anti-mouse Alexa Fluor 555 (Life Technologies), goat anti-mouse Alexa Fluor 488 (Life Technologies), goat anti-rabbit Alexa Fluor 488 (Life Technologies) and goat anti-chicken Alexa Fluor 488 (Life Technologies). Sections were imaged on an Olympus BX63F upright epifluorescence microscope using UPlanSAPO 10×/0.4, 20×/0.75, 40×/0.95, 60×/1.35 and 100×/1.35 objective lenses. H&E-stained slides were imaged on an Olympus VS120-L100-W Slide Scanner. Rabbit mammary tissue was collected and fixed in NBF (10%), processed and sectioned at 5 µm, as previously described *(64)*. Hematoxylin and eosin (H&E) staining was performed using standard histological protocols.

### CUBIC tissue clearing and wholemount immunostaining

Mammary tissue was dissected and fixed for 6-9 h, as previously optimized for wholemount immunostaining in the mammary gland *(9, 47)*. Tissue clearing was performed using a modified CUBIC protocol (CUBIC 1A) *(47, 65)* or SeeDB *(66)*. Wholemount immunostaining was performed following immersion in CUBIC Reagent 1A for 1-3 days (depending on tissue size) and overnight blocking in PBS with normal goat serum (10%) and triton X-100 (0.5%) *(47)*. SeeDB immersion clearing was performed as previously described *(3)*. The following primary antibodies were used in this study: rabbit SMA (1:300) and PGP9.5 (1:150). DAPI (5 μg/mL) staining was performed at room temperature for 2 h. For imaging CellTracker™ in cleared mammary tissue, tissue was loaded with CellTracker™ Red (1.5 μM) in complete media for 30 min prior to fixing. All tissue cleared using the CUBIC protocol was immersed in CUBIC Reagent 2 for at least 24 h prior to imaging. Cleared and immunostained tissue was imaged using an Olympus FV3000 laser scanning confocal microscope with UPLSAPO 10×/0.40, UPLSAPO 20×/0.75, UPLSAPO 30×/1.05 and UPLFLN 40×/0.75 objective lenses. Visualization and image processing was performed in ImageJ (v1.52e, National Institutes of Health) *(67, 68)*. Denoising of 3D image sequences was performed as previously described *(69)*.

### Wholemount histochemistry

Intact abdominal and inguinal mammary glands were dissected, spread on Tetra-Pak card and fixed overnight in 10% NBF at room temperature. Fixed tissue was immersed in CUBIC reagent 1 for 3 days at 37°C and placed in methyl green (0.5%) for 1.5 h. Following staining, tissue was rinsed twice in tap water and once in distilled water, prior to de-staining for 20 min in acid alcohol [50% ethanol and HCl (25 mM)]. After de-staining, tissue was transferred to CUBIC reagent 2 for 24 h and imaged using an Olympus SZX10 stereo microscope with a DF PLAPO 1× objective.

### Real-time RT-PCR

For RNA isolation from HC11 cell cultures, cells were lysed in QIAGEN Buffer RLT Plus. For RNA isolation from mouse mammary tissue, tissue was dissected from the abdominal mammary gland of virgin, lactating (day 10) and involuting (24 h) mice and immediately snap frozen in liquid nitrogen. Frozen tissue pieces (50-100 mg) were crushed using a mortar and pestle, resuspended in QIAGEN Buffer RLT Plus and homogenized using QIAshredder homogenizer columns (79651, QIAGEN). RNA was isolated and purified using the RNeasy Plus Mini Kit with gDNA Eliminator columns (74134, QIAGEN). Reverse transcription was performed using the QuantiTect Reverse Transcription Kit with integrated removal of genomic DNA contamination (205311, QIAGEN), and resulting cDNA was amplified using an Applied Biosystems ViiA 7 Real-time PCR System, using TaqMan Fast Universal PCR Master Mix (4352042, Applied Biosystems) and TaqMan Gene Expression Assays (Applied Biosystems). Relative quantification was calculated with reference to 18S ribosomal RNA and analysed using the comparative C_T_ method *(70)*.

### Immunoblotting

HC11 total cellular protein was isolated using protein lysis buffer supplemented with protease and phosphatase inhibitors (A32959, ThermoFisher), and immunoblotting was performed as previously described *(71)*. The following primary antibodies were used in this study: mouse β-actin (1:10,000), rabbit phospho-STAT5 (1:1000), mouse anti-STAT5 (1:500), and incubated at 4°C overnight. The following secondary antibodies were used: goat anti-mouse IgG HRP conjugate (1:10,000) and goat anti-rabbit IgG HRP conjugate (1:10,000), and incubated for 1 h at room temperature. Images were acquired using a Fusion SL chemiluminescence imaging system (Vilber Lourmat).

### Electron microscopy

For electron microscopy studies, animals were euthanized and abdominal and inguinal mammary glands were immediately harvested and processed into approx. 3 mm^3^ pieces in the presence of fixative. For serial block-face electron microscopy (SBEM), samples were processed as per Webb and Schieber *(72)*. Samples were imaged and sectioned using a Zeiss Sigma scanning electron microscope fitted with a 3View (Gatan). The resultant SBEM images were aligned using the StackReg ImageJ plugin *(73)* and image segmentation performed semi-automatically via the Ilastik segmentation toolkit carving workflow *(74)*. Image segmentation was further proofed and corrected using the Seg3D open-source software package *(75)*. SBEM movies were created in virtual reality using syGlass *(76)*.

For scanning electron microscopy (SEM), fixed samples were fractured and images acquired using a JCM-5000 NeoScope (Jeol).

### Basal cell contraction analysis

Basal cell surface area (SA) was analyzed using the surface tool in Imaris 9.2.1 (Bitplane). Cells were selected based on intensity and visibility in the field of view (5 cells per animal) and SA for each cell measured prior to oxytocin stimulation (range 3.25-19.54 s) and at maximal contraction post-oxytocin stimulation (range 104-472.27 s).

### Luminal cell size measurements

Luminal cell length (basal to apical, maximum distance), width (lateral, widest point) and area were calculated in FIJI using the straight line and freehand drawing tools and measure function.

### Histological analyses

Histological evaluation of mammary tissue was performed blinded by a veterinary pathologist (K.H.).

### Statistics

Statistical analysis was performed in GraphPad Prism (v7.03). Statistical tests are detailed in the figure legend. Standard error of the mean (S.E.M) is used throughout.

## Acknowledgments

The authors acknowledge the Translational Research Institute (TRI) for the research space, equipment and core facilities that enabled this research. We thank the UQ Biological Resource Facility staff for help with animal work; Dr Corinne Alberthsen for assistance with research ethics applications and compliance; Mr Richard Webb, Ms Robyn Chapman and Dr Kathryn Green (Centre for Microscopy and Microanalysis) for their support with the electron microscopy studies; Dr Jerome Boulanger (MRC Laboratory of Molecular Biology) for the 3D denoising algorithm; and Dr Adam Ewing and Mr Eric Pizzani (Translational Research Institute) for research computing support.

## Author contributions

Conceptualization, T.A.S. and F.M.D.; Methodology, T.A.S. and F.M.D.; Formal analysis, T.A.S., K.H., A.S., A.L.J. and F.M.D.; Investigation, T.A.S., A.S., K.H., and F.M.D.; Resources, N.M. (Komen Tissue Bank Samples); Writing – Original Draft, T.A.S. and F.M.D.; Visualization, T.A.S., M.M., A.L.J. and F.M.D.; Supervision, F.M.D.; Project Administration, F.M.D.; Funding Acquisition, F.M.D.

## Data and materials availability

All data needed to evaluate the conclusions in the paper are present in the paper and/or Supplementary Materials.

## Supplementary Materials

**Figure S1:**
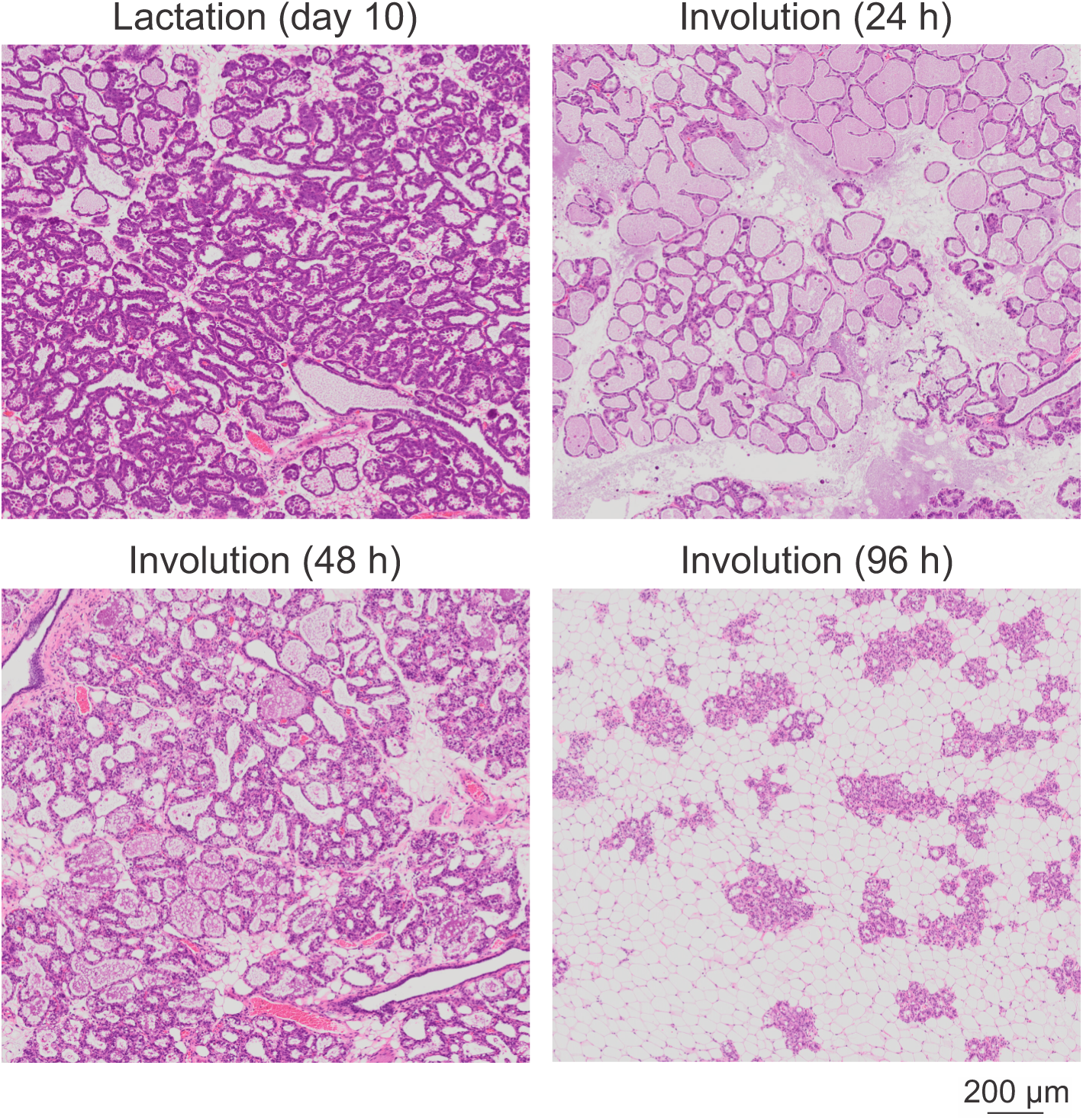
H&E staining of lactating (day 10) and involuting (24, 48, 96 h) mouse mammary tissue. Related to Figure 1. Representative of n = 3 mice.

**Figure S2:**
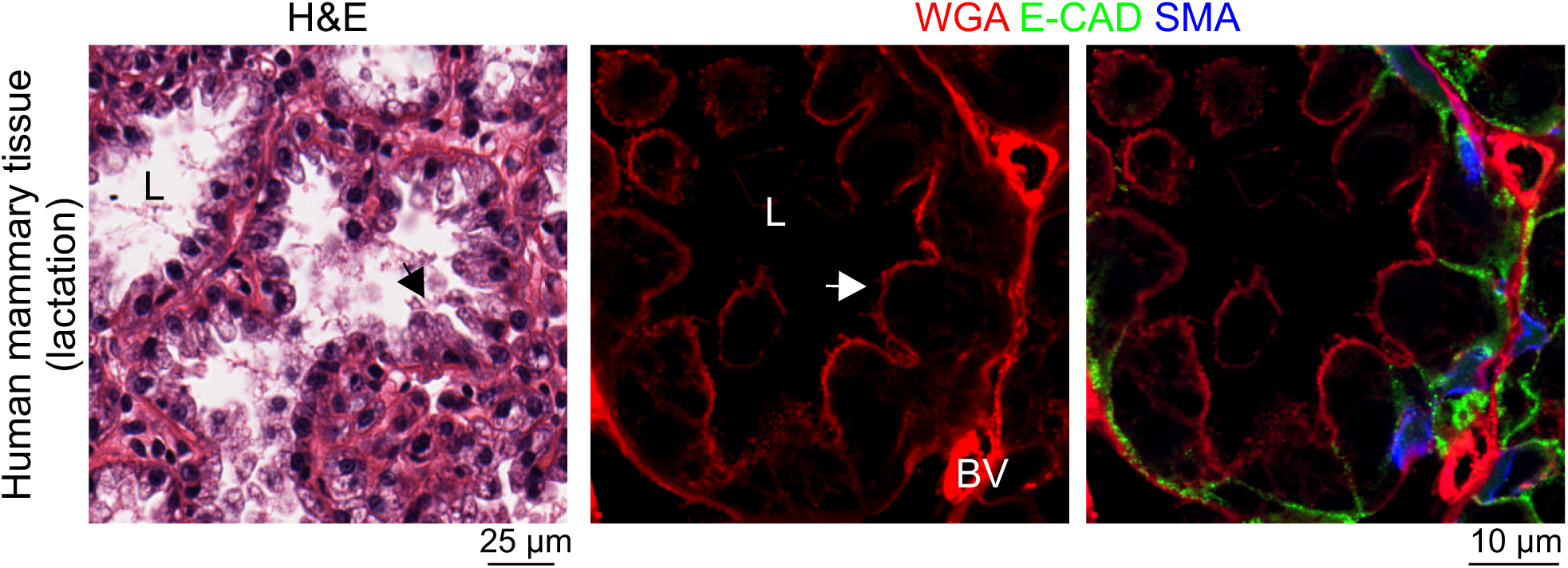
Human mammary tissue isolated during lactation. H&E (left) and fluorescence imaging (right) with WGA staining (red), E-cadherin immunostaining (green) and SMA immunostaining (blue). Arrow shows apical membrane curvature (beyond lateral contact sites). L, lumen; BV, blood vessel. Representative of n = 3 lactating human donors.

**Figure S3:**
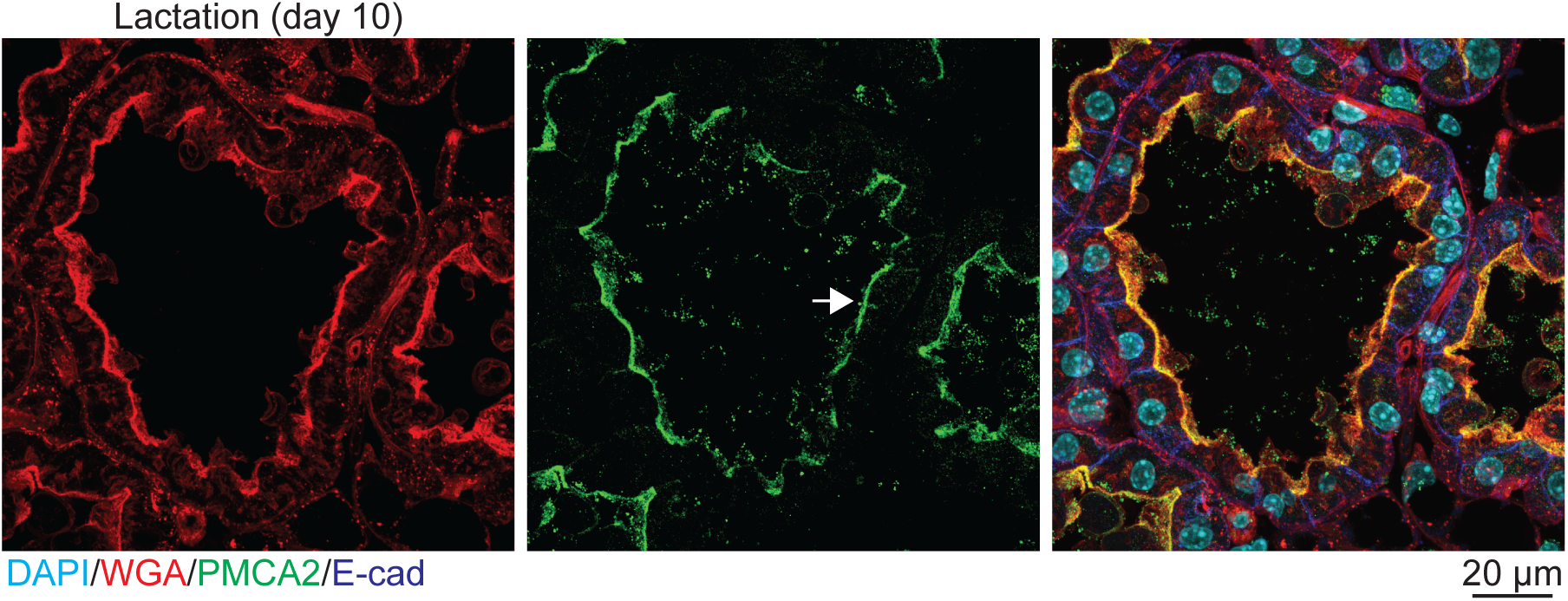
Immunofluorescence staining of mouse mammary tissue during lactation (day 10). WGA (red), PMCA2 (green), E-cadherin (blue), DAPI/nuclei (cyan). Arrow shows apical PMCA2 expression.

**Figure S4:**
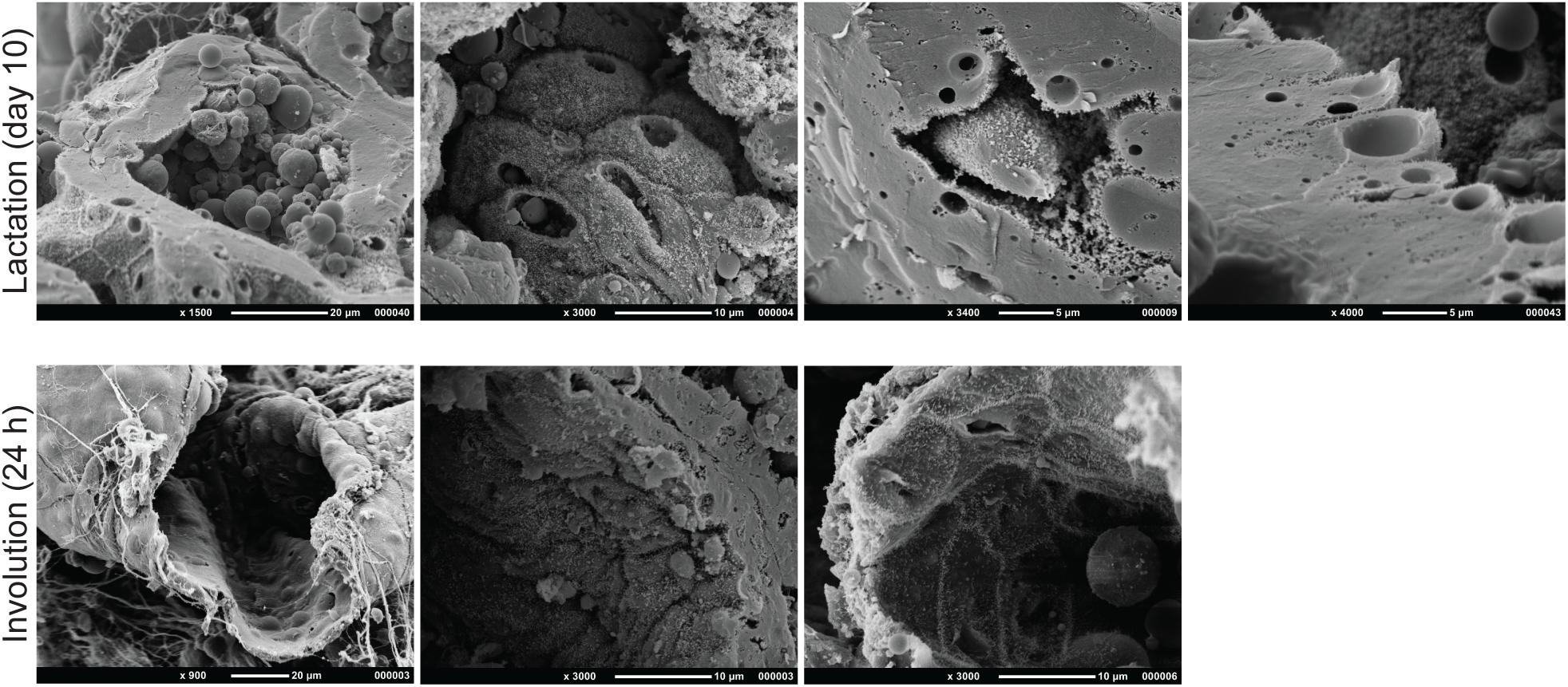
Additional SEM images of freeze-fractured alveoli during lactation and involution. Related to Figure 2D. Representative of n = 3 mice.

**Figure S5:**
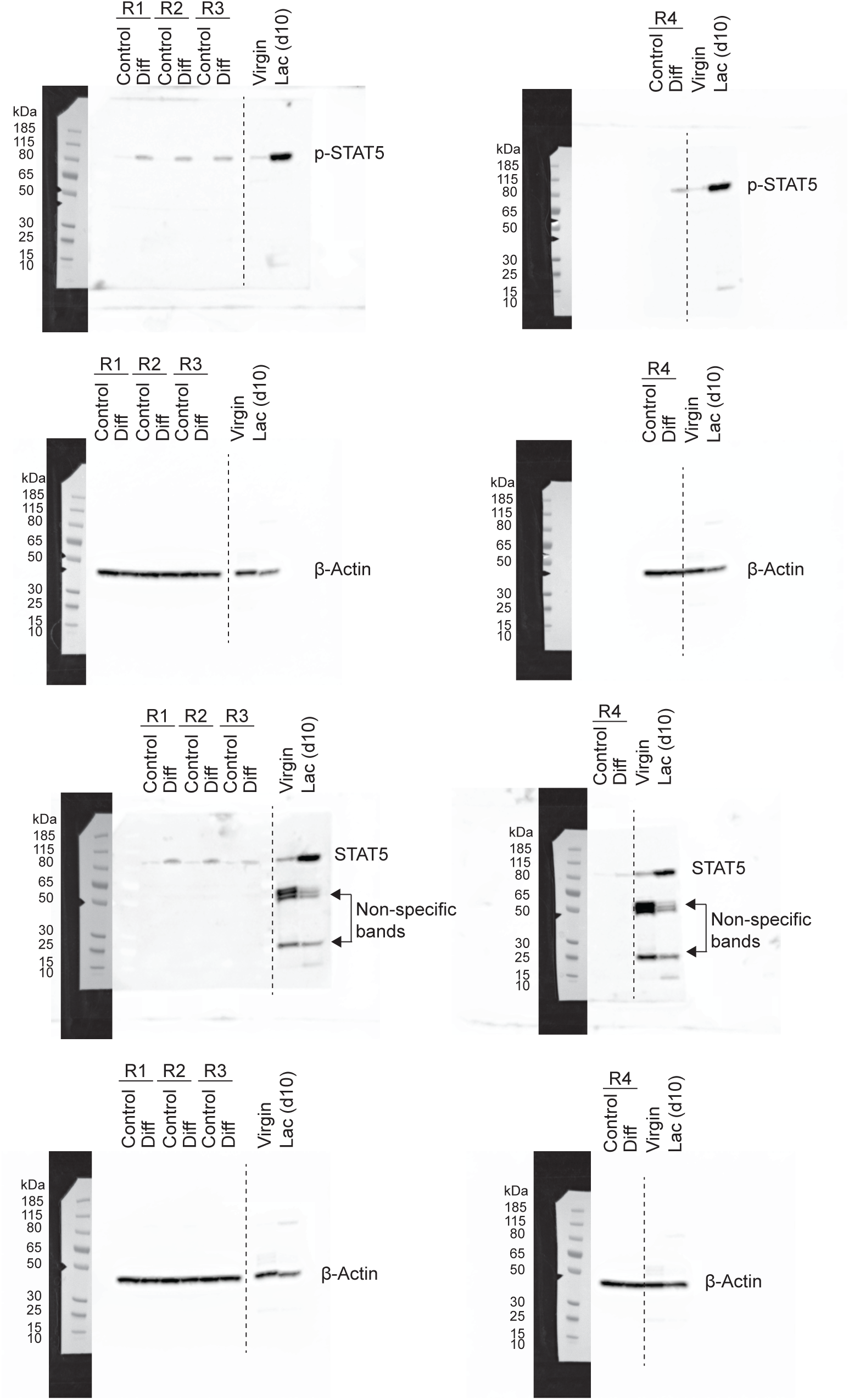
Full immunoblots related to Figure 3B.

**Figure S6:**
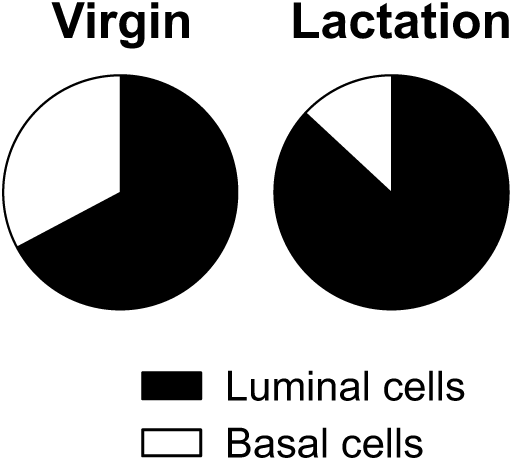
Proportion of luminal and basal cells (relative to total epithelial cell count) in mouse mammary tissue isolated from virgin and lactating mice; n = 3 mice.

**Figure S7:**
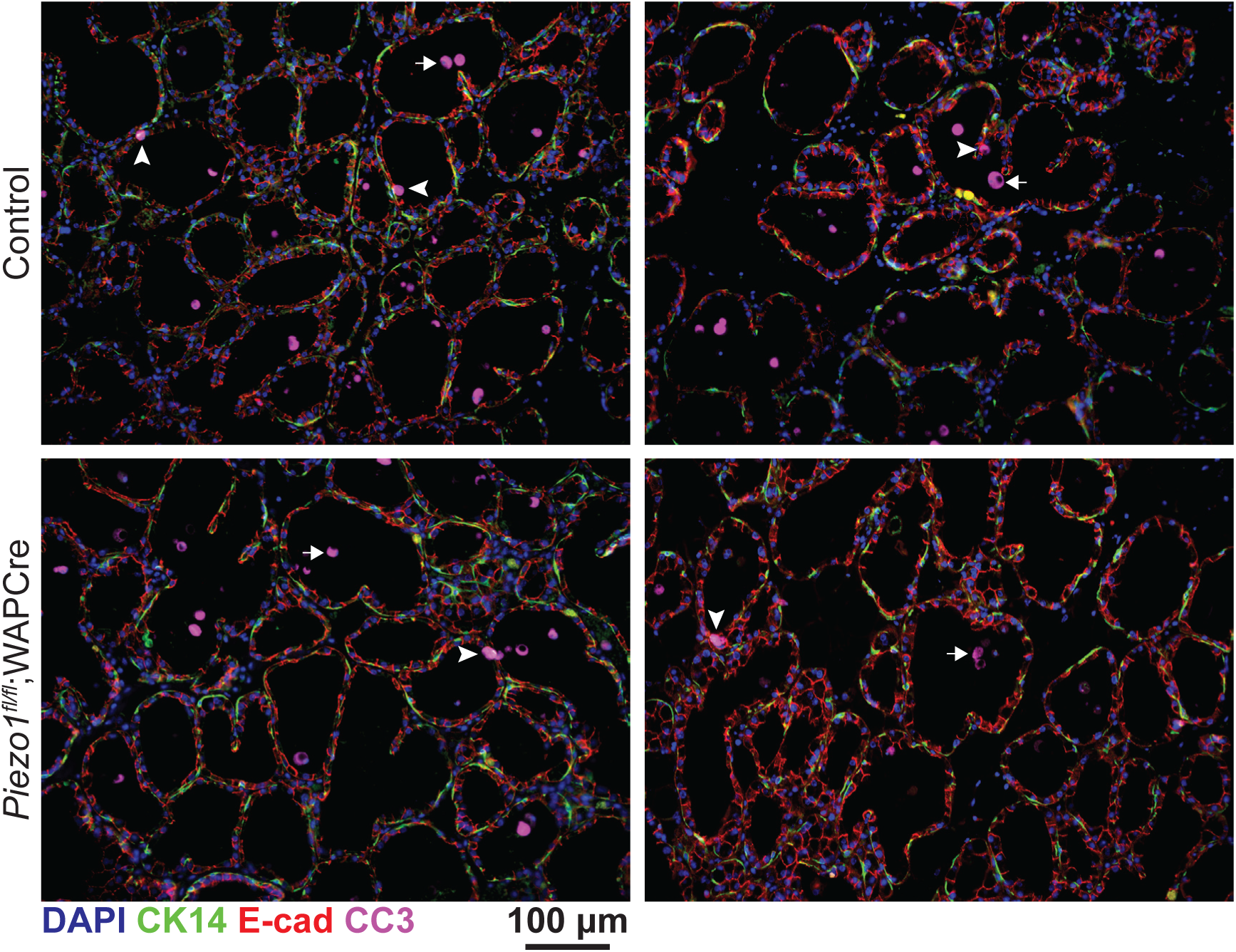
Immunofluorescence staining of mouse mammary tissue at 24 h after forced weaning for cleaved caspase 3 (CC3, magenta), CK14 (green) and E-cadherin (red). Nuclei are stained with DAPI (blue). CC3 positive cells were detected in both the alveolar lumen (arrow) and alveolar unit wall (arrow head). Related to Figure 5. Images have been brightness adjusted for display purposes.

**Figure S8:**
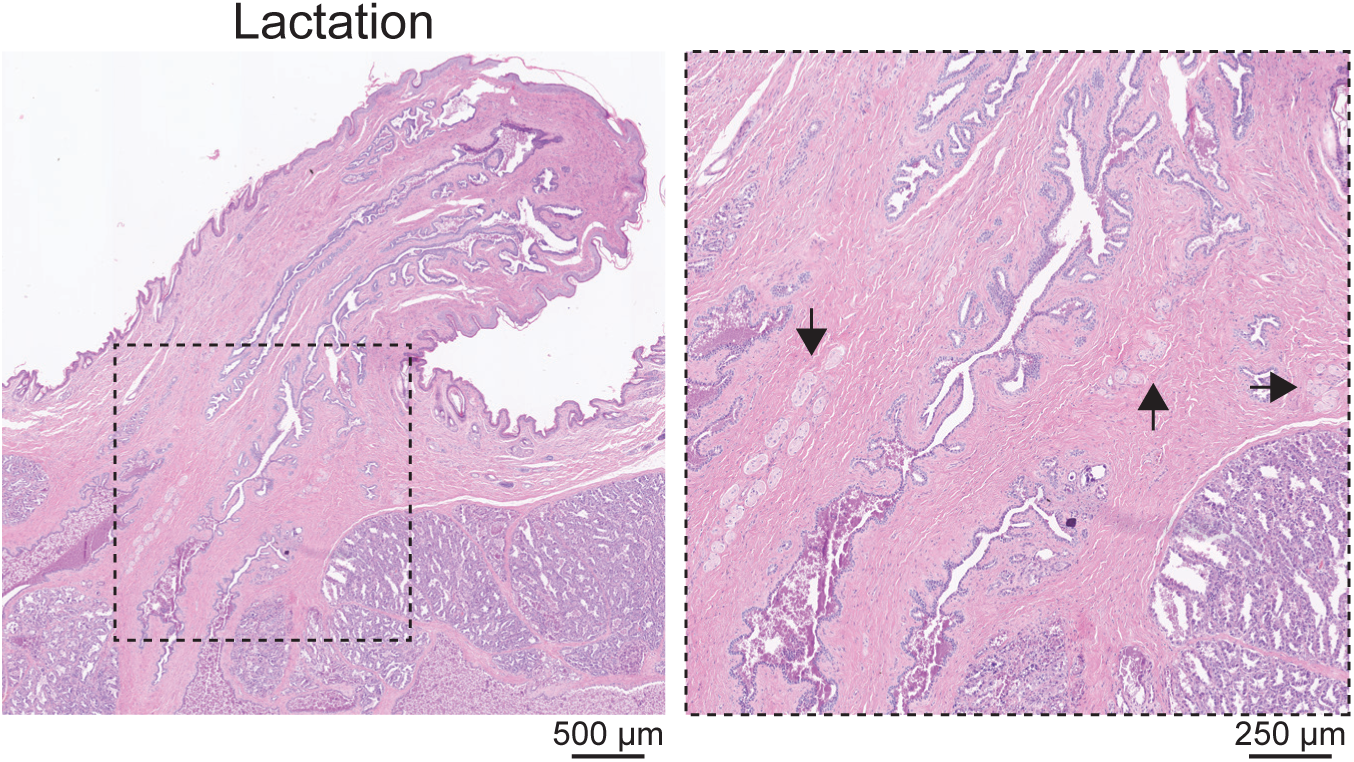
H&E staining of lactating rabbit teat and mammary tissue. Boxed region is magnified in adjacent sub-panel; arrows show neural tissue. Related to Figure 6. Representative of n = 3 rabbits.

Movie S1. Depth-coded movie showing live lactating mammary tissue stained with CellTracker™ Red and stimulated with oxytocin (85 nM) approximately 3-5 minutes before imaging. Total movie length is 16 min 30 s.

Movie S2. SBEM imaging of lactating mouse mammary tissue (no rendering). Movie created from 999 × 50 nm sections.

Movie S3. SBEM imaging of lactating mouse mammary tissue. Rendering highlights luminal epithelial cells (purple, pink and dark green), nucleus (light green) and multiple milk fat globules (round objects, multi colored). Movie shows 350 × 50 nm sections.

Movie S4. SBEM imaging of involuting (24 h) mouse mammary tissue (no rendering). Movie shows 1000 × 50 nm sections.

Movie S5. SBEM imaging of involuting (24 h) mouse mammary tissue. Rendering highlights luminal epithelial cell (blue), milk fat globule (yellow) and nucleus (green). Movie shows 280 × 50 nm sections.

Movie S6. Control (left) and differentiated (right) HC11 mouse mammary epithelial cells loaded with the ratiometric calcium indicator Fura-5F. Recordings show calcium responses (340/380) in pseudocolor to fluid shear stress. Total movie length is 150 s. Montage shows 9 individual wells for each condition (labelled in first frame) from 3 independent experiments.

Movie S7. Primary cells isolated from pregnant *GCaMP6f;K8CreERT2* mice and loaded with CellTracker™ Red. Cells were treated with vehicle (physiological salt solution) at frame 10 (30 s). Total movie length is 6 min.

Movie S8. Primary cells isolated from pregnant *GCaMP6f;K8CreERT2* mice and loaded with CellTracker™ Red. Cells were treated with Yoda1 (12 µM) in physiological salt solution at frame 10 (30 s). Total movie length is 6 min.

Movie S9. Primary cells isolated from pregnant *GCaMP6f-TdTom;K5CreERT2* mice. Cells were treated with vehicle (physiological salt solution) at frame 10 (30 s). Cells were stimulated with oxytocin at the end of the assay to confirm their responsiveness. Total movie length is 6 min 27 s.

Movie 10. Primary cells isolated from pregnant *GCaMP6f-TdTom;K5CreERT2* mice. Cells were treated with Yoda1 (12 µM) in physiological salt solution at frame 10 (30 s). Cells were stimulated with oxytocin at the end of the assay to confirm their responsiveness. Total movie length is 6 min 27 s.

## References

1. H. Macias, L. Hinck, Mammary gland development, Wiley Interdiscip. Rev. Dev. Biol. 1, 533–557 (2012).

2. A. Van Keymeulen, A. S. Rocha, M. Ousset, B. Beck, G. Bouvencourt, J. Rock, N. Sharma, S. Dekoninck, C. Blanpain, Distinct stem cells contribute to mammary gland development and maintenance, Nature 479, 189–193 (2011).

3. F. M. Davis, B. Lloyd-Lewis, O. B. Harris, S. Kozar, D. J. Winton, L. Muresan, C. J. Watson, Single-cell lineage tracing in the mammary gland reveals stochastic clonal dispersion of stem/progenitor cell progeny, Nat. Commun. 7, 13053 (2016).

4. A. Van Keymeulen, M. Fioramonti, A. Centonze, G. Bouvencourt, Y. Achouri, C. Blanpain, Lineage-restricted mammary stem cells sustain the development, homeostasis, and regeneration of the estrogen receptor positive lineage, Cell Rep. 20, 1525–1532 (2017).

5. C. Scheele, E. Hannezo, M. Muraro, A. Zomer, N. Langedijk, A. van Oudenaarden, B. Simons, J. van Rheenen, Identity and dynamics of mammary stem cells during branching morphogenesis, Nature (2017).

6. A. Sreekumar, M. J. Toneff, E. Toh, K. Roarty, C. J. Creighton, G. K. Belka, D.-K. Lee, J. Xu, L. A. Chodosh, J. S. Richards, J. M. Rosen, WNT-mediated regulation of FOXO1 constitutes a critical axis maintaining pubertal mammary stem cell homeostasis, Dev. Cell (2017).

7. B. Lloyd-Lewis, O. B. Harris, C. J. Watson, F. M. Davis, Mammary stem cells: Premise, properties and perspectives, Trends Cell Biol. 8, 556–567 (2017).

8. R. K. Zwick, M. C. Rudolph, B. A. Shook, B. Holtrup, E. Roth, V. Lei, A. Van Keymeulen, V. Seewaldt, S. Kwei, J. Wysolmerski, M. S. Rodeheffer, V. Horsley, Adipocyte hypertrophy and lipid dynamics underlie mammary gland remodeling after lactation, Nat. Commun. (2018).

9. B. Lloyd-Lewis, F. M. Davis, O. B. Harris, J. R. Hitchcock, C. J. Watson, Neutral lineage tracing of proliferative embryonic and adult mammary stem/progenitor cells, Development 145, dev164079 (2018).

10. J. L. McManaman, M. C. Neville, Mammary physiology and milk secretionAdv. Drug Deliv. Rev. (2003).

11. F. M. Davis, The ins and outs of calcium signalling in lactation and involution: implications for breast cancer treatment, Pharmacol. Res. 116, 100–104 (2016).

12. F. M. Davis, A. Janoshazi, K. S. Janardhan, N. Steinckwich, D. M. D’Agostin, J. G. Petranka, P. N. Desai, S. J. Roberts-Thomson, G. S. Bird, D. K. Tucker, S. E. Fenton, S. Feske, G. R. Monteith, J. W. Putney, Essential role of Orai1 store-operated calcium channels in lactation, Proc. Natl. Acad. Sci. 112, 5827–5832 (2015).

13. T. J. Sargeant, B. Lloyd-Lewis, H. K. Resemann, A. Ramos-Montoya, J. Skepper, C. J. Watson, Stat3 controls cell death during mammary gland involution by regulating uptake of milk fat globules and lysosomal membrane permeabilization, Nat. Cell Biol. 16, 1057–68 (2014).

14. P. A. Kreuzaler, A. D. Staniszewska, W. Li, N. Omidvar, B. Kedjouar, J. Turkson, V. Poli, R. A. Flavell, R. W. E. Clarkson, C. J. Watson, Stat3 controls lysosomal-mediated cell death in vivo, Nat. Cell Biol. 13, 303–9 (2011).

15. C. J. Watson, P. A. Kreuzaler, Remodeling mechanisms of the mammary gland during involution, Int. J. Dev. Biol. 55, 757–762 (2011).

16. C. Brisken, D. Ataca, Endocrine hormones and local signals during the development of the mouse mammary gland, Wiley Interdiscip. Rev. Dev. Biol. 4, 181–195 (2015).

17. G. Gimpl, F. Fahrenholz, The oxytocin receptor system: structure, function, and regulation, Physiol. Rev. 81, 629–683 (2001).

18. E. A. Kritikou, A. Sharkey, K. Abell, P. J. Came, E. Anderson, R. W. E. Clarkson, C. J. Watson, A dual, non-redundant, role for LIF as a regulator of development and STAT3-mediated cell death in mammary gland, Development 130, 3459–68 (2003).

19. K. Raymond, S. Cagnet, M. Kreft, H. Janssen, A. Sonnenberg, M. A. Glukhova, Control of mammary myoepithelial cell contractile function by α3β1 integrin signalling, EMBO J. 30, 1896–1906 (2011).

20. J. VanHouten, C. Sullivan, C. Bazinet, T. Ryoo, R. Camp, D. L. Rimm, G. Chung, J. Wysolmerski, PMCA2 regulates apoptosis during mammary gland involution and predicts outcome in breast cancer, Proc. Natl. Acad. Sci. U. S. A. 107, 11405–11410 (2010).

21. P. P. Provenzano, D. R. Inman, K. W. Eliceiri, P. J. Keely, Matrix density-induced mechanoregulation of breast cell phenotype, signaling and gene expression through a FAK-ERK linkage, Oncogene (2009).

22. C. Bonnans, J. Chou, Z. Werb, Remodelling the extracellular matrix in development and disease*Nat. Rev*. Mol. Cell Biol. (2014).

23. A. Quaglino, M. Salierno, J. Pellegrotti, N. Rubinstein, E. C. Kordon, Mechanical strain induces involution-associated events in mammary epithelial cells, BMC Cell Biol. (2009).

24. G. T. Eisenhoffer, P. D. Loftus, M. Yoshigi, H. Otsuna, C. Bin Chien, P. A. Morcos, J. Rosenblatt, Crowding induces live cell extrusion to maintain homeostatic cell numbers in epithelia, Nature 484, 546–549 (2012).

25. B. Coste, J. Mathur, M. Schmidt, T. J. Earley, S. Ranade, M. J. Petrus, A. E. Dubin, A. Patapoutian, Piezo1 and Piezo2 are essential components of distinct mechanically activated cation channels, Science (80-.). (2010).

26. S. A. Gudipaty, J. Lindblom, P. D. Loftus, M. J. Redd, K. Edes, C. F. Davey, V. Krishnegowda, J. Rosenblatt, Mechanical stretch triggers rapid epithelial cell division through Piezo1, Nature 543, 118–121 (2017).

27. K. Nonomura, V. Lukacs, D. Sweet, L. Goddard, A. Kanie, T. Whitwam, S. Ranade, T. Fujimori, M. Kahn, A. Patapoutian, Mechanically activated ion channel PIEZO1 is required for lymphatic valve formation, Proc. Natl. Acad. Sci. 115, 12817–12822 (2018).

28. S. M. Cahalan, V. Lukacs, S. S. Ranade, S. Chien, M. Bandell, A. Patapoutian, Piezo1 links mechanical forces to red blood cell volume, Elife (2015).

29. M. Tsuchiya, Y. Hara, M. Okuda, K. Itoh, R. Nishioka, A. Shiomi, K. Nagao, M. Mori, Y. Mori, J. Ikenouchi, R. Suzuki, M. Tanaka, T. Ohwada, J. Aoki, M. Kanagawa, T. Toda, Y. Nagata, R. Matsuda, Y. Takayama, M. Tominaga, M. Umeda, Cell surface flip-flop of phosphatidylserine is critical for PIEZO1-mediated myotube formation, Nat. Commun. (2018).

30. J. M. J. Romac, R. A. Shahid, S. M. Swain, S. R. Vigna, R. A. Liddle, Piezo1 is a mechanically activated ion channel and mediates pressure induced pancreatitis, Nat. Commun. (2018).

31. C. J. Haaksma, R. J. Schwartz, J. J. Tomasek, Myoepithelial cell contraction and milk ejection are impaired in mammary glands of mice lacking smooth muscle alpha-actin, Biol. Reprod. 85, 13–21 (2011).

32. C. Stevens, P. J. Hunter, in Progress in Biophysics and Molecular Biology, (2003).

33. Y. Tokita, H. Akiho, K. Nakamura, E. Ihara, M. Yamamoto, Contraction of gut smooth muscle cells assessed by fluorescence imaging, J. Pharmacol. Sci. (2015).

34. K. Hughes, C. J. Watson, C. J. Watson, The role of Stat3 in mammary gland involution: Cell death regulator and modulator of inflammation, Horm. Mol. Biol. Clin. Investig. (2012).

35. M. Li, X. Liu, G. Robinson, U. Bar-Peled, K.-U. Wagner, W. S. Young, L. Hennighausen, P. A. Furth, Mammary-derived signals activate programmed cell death during the first stage of mammary gland involution, Proc. Natl. Acad. Sci. (2002).

36. K. Hughes, J. A. Wickenden, J. E. Allen, C. J. Watson, Conditional deletion of Stat3 in mammary epithelium impairs the acute phase response and modulates immune cell numbers during post-lactational regression, J. Pathol. (2012).

37. W. Doppler, B. Groner, R. K. Ball, Prolactin and glucocorticoid hormones synergistically induce expression of transfected rat beta-casein gene promoter constructs in a mammary epithelial cell line., Proc. Natl. Acad. Sci. (1989).

38. J. Xing, J. G. J. G. Petranka, F. M. F. M. Davis, P. N. P. N. Desai, J. W. J. W. Putney, G. S. G. S. Bird, Role of Orai1 and store-operated calcium entry in mouse lacrimal gland signalling and function, J. Physiol. 592, 927–39 (2014).

39. J. Xu, J. Mathur, E. Vessières, S. Hammack, K. Nonomura, J. Favre, L. Grimaud, M. Petrus, A. Francisco, J. Li, V. Lee, F. Xiang, J. Mainquist, S. Cahalan, A. Orth, J. Walker, S. Ma, V. Lukacs, L. Bordone, M. Bandell, B. Laffitte, Y. Xu, S. Chien, D. Henrion, A. Patapoutian, GRP68 senses flow and is essential for vascular physiology, Cell 173, 762–75 (2018).

40. B. Martinac, The ion channels to cytoskeleton connection as potential mechanism of mechanosensitivity*Biochim*. Biophys. Acta - Biomembr. (2014).

41. K. Bach, S. Pensa, M. Grzelak, J. Hadfield, D. J. Adams, J. C. Marioni, W. T. Khaled, Differentiation dynamics of mammary epithelial cells revealed by single-cell RNA sequencing, Nat. Commun. 8 (2017).

42. T. A. Reinhardt, R. L. Horst, Ca2+-ATPases and their expression in the mammary gland of pregnant and lactating rats, Am. J. Physiol. 276, C796–C802 (1999).

43. D. McAndrew, D. M. D. M. Grice, A. A. A. A. Peters, F. M. F. M. Davis, T. Stewart, M. Rice, C. E. C. E. Smart, M. A. M. A. Brown, P. A. P. A. Kenny, S. J. S. J. Roberts-Thomson, G. R. G. R. Monteith, ORAI1-mediated calcium influx in lactation and in breast cancer, Mol. Cancer Ther. 10, 448–460 (2011).

44. R. Syeda, J. Xu, A. E. Dubin, B. Coste, J. Mathur, T. Huynh, J. Matzen, J. Lao, D. C. Tully, I. H. Engels, H. Michael Petrassi, A. M. Schumacher, M. Montal, M. Bandell, A. Patapoutian, Chemical activation of the mechanotransduction channel Piezo1, Elife 4, e07369 (2015).

45. T.-W. Chen, T. J. Wardill, Y. Sun, S. R. Pulver, S. L. Renninger, A. Baohan, E. R. Schreiter, R. A. Kerr, M. B. Orger, V. Jayaraman, L. L. Looger, K. Svoboda, D. S. Kim, Ultrasensitive fluorescent proteins for imaging neuronal activity., Nature 499, 295–300 (2013).

46. K. U. Wagner, R. J. Wall, L. St-Onge, P. Gruss, A. Wynshaw-Boris, L. Garrett, M. Li, P. A. Furth, L. Hennighausen, Cre-mediated gene deletion in the mammary gland, Nucleic Acids Res. 25, 4323–4330 (1997).

47. B. Lloyd-Lewis, F. M. Davis, O. B. Harris, J. R. Hitchcock, F. C. Lourenco, M. Pasche, C. J. Watson, Imaging the mammary gland and mammary tumours in 3D: Optical tissue clearing and immunofluorescence methods, Breast Cancer Res. 18 (2016).

48. B. Lloyd-Lewis, T. J. Sargeant, P. A. Kreuzaler, H. K. Resemann, S. Pensa, C. J. Watson, in Methods in Molecular Biology, (2017).

49. V. Djonov, A. C. Andres, A. Ziemiecki, Vascular remodelling during the normal and malignant life cycle of the mammary gland*Microsc*. Res. Tech. 52, 182–189 (2001).

50. S. S. Ranade, Z. Qiu, S.-H. Woo, S. S. Hur, S. E. Murthy, S. M. Cahalan, J. Xu, J. Mathur, M. Bandell, B. Coste, Y.-S. J. Li, S. Chien, A. Patapoutian, Piezo1, a mechanically activated ion channel, is required for vascular development in mice, Proc. Natl. Acad. Sci. 111, 10347–10352 (2014).

51. S. S. Ranade, S. H. Woo, A. E. Dubin, R. A. Moshourab, C. Wetzel, M. Petrus, J. Mathur, V. Bégay, B. Coste, J. Mainquist, A. J. Wilson, A. G. Francisco, K. Reddy, Z. Qiu, J. N. Wood, G. R. Lewin, A. Patapoutian, Piezo2 is the major transducer of mechanical forces for touch sensation in mice, Nature (2014).

52. M. Bringmann, D. C. Bergmann, Tissue-wide Mechanical Forces Influence the Polarity of Stomatal Stem Cells in Arabidopsis, Curr. Biol. (2017).

53. J. C. Tuthill, R. I. Wilson, Mechanosensation and Adaptive Motor Control in Insects*Curr*. Biol. (2016).

54. B. Pan, N. Akyuz, X. P. Liu, Y. Asai, C. Nist-Lund, K. Kurima, B. H. Derfler, B. György, W. Limapichat, S. Walujkar, L. N. Wimalasena, M. Sotomayor, D. P. Corey, J. R. Holt, TMC1 Forms the Pore of Mechanosensory Transduction Channels in Vertebrate Inner Ear Hair Cells, Neuron (2018).

55. S. H. Woo, V. Lukacs, J. C. De Nooij, D. Zaytseva, C. R. Criddle, A. Francisco, T. M. Jessell, K. A. Wilkinson, A. Patapoutian, Piezo2 is the principal mechanotransduction channel for proprioception, Nat. Neurosci. (2015).

56. W. Z. Zeng, K. L. Marshall, S. Min, I. Daou, M. W. Chapleau, F. M. Abboud, S. D. Liberles, A. Patapoutian, PIEZOs mediate neuronal sensing of blood pressure and the baroreceptor reflex, Science (80-.). (2018).

57. J. Raeburn, Release of oxytocin and prolactin in response to suckling, BMJ (2009).

58. A. C. Rios, N. Y. Fu, G. J. Lindeman, J. E. Visvader, In situ identification of bipotent stem cells in the mammary gland, Nature 506, 322–7 (2014).

59. N. E. Hynes, D. Taverna, I. M. Harwerth, F. Ciardiello, D. S. Salomon, T. Yamamoto, B. Groner, Epidermal growth factor receptor, but not c-erbB-2, activation prevents lactogenic hormone induction of the beta-casein gene in mouse mammary epithelial cells., Mol. Cell. Biol. (2015).

60. M. E. Sherman, J. D. Figueroa, J. E. Henry, S. E. Clare, C. Rufenbarger, A. M. Storniolo, The Susan G. Komen for the Cure Tissue Bank at the IU Simon Cancer Center: A unique resource for defining the “molecular histology” of the breast, Cancer Prev. Res. 5, 528–535 (2012).

61. J. J. Bassett, A. H. L. Bong, E. K. Janke, M. Robitaille, S. J. Roberts-Thomson, A. A. Peters, G. R. Monteith, Assessment of cytosolic free calcium changes during ceramide-induced cell death in MDA-MB-231 breast cancer cells expressing the calcium sensor GCaMP6m, Cell Calcium (2018).

62. R. Rudolf, P. J. Magalhães, T. Pozzan, Direct in vivo monitoring of sarcoplasmic reticulum Ca2+ and cytosolic cAMP dynamics in mouse skeletal muscle., J. Cell Biol. 173, 187–93 (2006).

63. M. Prater, M. Shehata, C. J. Watson, J. Stingl, Enzymatic dissociation, flow cytometric analysis, and culture of normal mouse mammary tissue, Methods Mol.Biol. 946, 395–409 (2013).

64. K. Hughes, C. J. Watson, Sinus-like dilatations of the mammary milk ducts, Ki67 expression, and CD3-positive T lymphocyte infiltration, in the mammary gland of wild European rabbits during pregnancy and lactation, J. Anat. (2018).

65. E. A. Susaki, H. R. Ueda, Whole-body and Whole-Organ Clearing and Imaging Techniques with Single-Cell Resolution: Toward Organism-Level Systems Biology in Mammals, Cell Chem. Biol. 23, 137–157 (2016).

66. M.-T. Ke, S. Fujimoto, T. Imai, SeeDB: a simple and morphology-preserving optical clearing agent for neuronal circuit reconstruction., Nat. Neurosci. 16, 1154–61 (2013).

67. J. Schindelin, I. Arganda-Carreras, E. Frise, V. Kaynig, M. Longair, T. Pietzsch, S. Preibisch, C. Rueden, S. Saalfeld, B. Schmid, J.-Y. J.-Y. Tinevez, D. J. White, V. Hartenstein, K. Eliceiri, P. Tomancak, A. Cardona, K. Liceiri, P. Tomancak, C. A., Fiji: an open source platform for biological image analysis., Nat. Methods 9, 676–682 (2012).

68. M. Linkert, C. T. Rueden, C. Allan, J. M. Burel, W. Moore, A. Patterson, B. Loranger, J. Moore, C. Neves, D. MacDonald, A. Tarkowska, C. Sticco, E. Hill, M. Rossner, K. W. Eliceiri, J. R. Swedlow, Metadata matters: access to image data in the real world J. Cell Biol. 189, 777–782 (2010).

69. J. Boulanger, C. Kervrann, P. Bouthemy, P. Elbau, J.-B. Sibarita, J. Salamero, Patch-based nonlocal functional for denoising fluorescence microscopy image sequences., IEEE Trans. Med. Imaging 29, 442–54 (2010).

70. K. M. Suchanek, F. J. May, J. A. Robinson, W. J. Lee, N. A. Holman, G. R. Monteith, S. J. Roberts-Thomson, Peroxisome proliferator-activated receptor alpha in the human breast cancer cell lines MCF-7 and MDA-MB-231, Mol. Carcinog. 34, 165–171 (2002).

71. F. M. Davis, I. Azimi, R. A. Faville, A. A. Peters, K. Jalink, J. W. Putney, G. J. Goodhill, E. W. Thompson, S. J. Roberts-Thomson, G. R. Monteith, Induction of epithelial-mesenchymal transition (EMT) in breast cancer cells is calcium signal dependent, Oncogene 33, 2307–2316 (2014).

72. R. Webb, N. Schieber, in Cellular Imaging. Biological and Medical Physics, Biomedical Engineering., E. Hanssen, Ed. (Springer, Cham, 2018), pp. 117–148.

73. P. Thévenaz, U. E. Ruttimann, M. Unser, A pyramid approach to subpixel registration based on intensity, IEEE Trans. Image Process. (1998).

74. C. Sommer, C. Straehle, U. Kothe, F. A. Hamprecht, in Proceedings - International Symposium on Biomedical Imaging, (2011).

75. CIBC, Seg3DVol. Image Segmentation Vis. Sci. Comput. Imaging Inst. (2015).

76. S. Pidhorskyi, M. Morehead, Q. Jones, G. Spirou, G. Doretto, syGlass: Interactive Exploration of Multidimensional Images Using Virtual Reality Head-mounted Displays (2018).

